# Controlled Delivery of a Neurotrophic Factor in the Adult Mouse Brain Using Engineered Microglia

**DOI:** 10.1101/2025.10.06.680702

**Authors:** Rohan J. Hofland, Marta Grońska-Pęski, Hiroko Nobuta, Nicolas Buitrago, Karan Malhotra, Jean M. Hébert, J. Tiago Gonçalves

## Abstract

Microglia, the resident immune cells of the central nervous system, have been proposed as vehicles for delivering therapeutic biologics. These cells can be genetically engineered in vitro and transplanted into host animals following ablation of endogenous microglia, enabling repopulation of the brain parenchyma. However, current replacement strategies often rely on radiation or transgenic models, limiting their clinical relevance. CSF1R inhibitors offer a more translational approach to microglia ablation, though surviving host cells can compete with transplanted microglia during repopulation.

In this study, we successfully ablated endogenous microglia using a CSF1R inhibitor in adult mice and developed a method to transplant engineered microglia expressing Brain-Derived Neurotrophic Factor (BDNF) in a doxycycline-inducible manner. To enhance engraftment, transplanted cells also expressed a constitutively active CSF1R mutant (caCSF1R).

BDNF-expressing transplanted microglia spread through large areas of host mice brains, displayed similar morphology and transcriptional profile to repopulating host microglia, and responded to pro-inflammatory stimuli. Treatment with doxycycline resulted in increased BDNF expression and TrkB phosphorylation in the host brain. Expression of caCSF1R provided transplanted cells with a competitive advantage over endogenous repopulating cells, resulting in the accelerated spread of the transplants.

Our results demonstrate the functional integration and therapeutic potential of microglia as vehicles for delivering neurotrophic factors to the brain in a controllable manner. Furthermore, we show that caCSF1R expression is able to enhance the spread of transplanted microglia.

**SIGNIFICANCE:** This study demonstrates the potential of engineered microglia to deliver the protein Brain-Derived Neurotrophic Factor to the brain parenchyma, under the control of orally-administered doxycycline. The technique can be generalized to a wide array of proteins, offering a novel paradigm for neurological therapy.

## INTRODUCTION

Biologics, biologically derived products, such as monoclonal antibodies and chimeric antigen receptor (CAR) T-cells, significantly improve outcomes for diseases like autoimmune disorders and cancers[1]. Due to the limited permeability of the Blood-Brain Barrier (BBB)[2, 3], current research on biologics for conditions such as stroke and Alzheimer’s disease faces a significant challenge in obtaining widespread distribution of the biologic throughout the brain. This prevents proper drug delivery and lowers the efficacy of the administered therapies.

Microglia, the immune cells of the CNS, are promising delivery vehicles for therapeutic biologics because of their unique ability to proliferate and retile entire brain regions after depletion[4–6]. Primary mouse microglia directly implanted in the adult mouse brain parenchyma, devoid of endogenous microglia, are able to successfully recolonize the brain [7], further supporting the idea of replacing endogenous microglia with engineered versions capable of delivering biologic drugs[8]. Depletion of host microglia is a critical step in microglial replacement, as microglia maintain homeostatic density through contact inhibition [9]. Once the treatment for depletion of microglia is stopped the brain is repopulated by a mix of surviving endogenous microglia (E-MG) [10] and, under some circumstances, infiltrating peripheral macrophages[11]. These repopulating microglia-like cells (R-MGLs), inhibit the spread of transplanted microglia (T-MG). To avoid this problem, many microglia depletion methods inhibit repopulation by R-MGLs using chemotherapy and/or full-body irradiation [7, 12, 13], which are clinically undesirable to the patients. Also, microglial engraftments can take a significant amount of time to repopulate the brain[14, 15]; thus, a more rapid repopulation of T-MG would reduce the time spent under microglia depletion and improve patient safety.

Neurotrophic factors are biomolecules with therapeutic relevance for neurodegenerative diseases. However, their size limits their efficient delivery across the blood-brain barrier, using systemic delivery methods. In this study, we transplanted primary microglia that were engineered to express Brain-Derived Neurotrophic Factor (BDNF), a growth factor deficient in AD patients[16, 17]. Transplanted microglia were also engineered to express a constitutively active mutant of the Colony Stimulating Factor 1 Receptor (caCSF1R), which gives them a competitive edge over R-MGLs in repopulating the brain following microglia ablation. Expression of both BDNF and caCSF1R was induced by doxycycline, allowing their expression to be regulated after transplantation. We show that T-MGs expressing caCSF1R repopulated the entirety of the hippocampus and the majority of the cortex 3 weeks post-transplantation, 3 times as fast as controls, and released biochemically functional BDNF in their vicinity. Analyses of the properties of early-stage T-MGs and R-MGLs showed that these two cell cohorts had similar transcriptional profiles, indicating a high degree of similarity between endogenous and transplanted repopulating cells. Our approach allowed for doxycycline-dependent BDNF delivery in wild-type mice pre-treated with a CSF1R antagonist drug, avoiding harmful methods like irradiation or chemotherapy[9, 13]. Our findings underscore the potential for customizable microglia-based delivery systems in CNS therapies.

## RESULTS

### Transplanted microglia engraft in a CNS environment devoid of endogenous microglia

To easily trace T-MGs in host animals, we developed a strategy to isolate and culture highly purified cortical T-MGs from *Cx3cr1^GFP^;Rosa26-rtTA* E15-P3 mouse pups, which express GFP under the *Cx3cr1* promoter, a monocyte marker primarily expressed by CNS microglia[18] (Supplemental figure 1A). By exploiting the adherent nature of T-MGs, we generated monolayer cultures of GFP+ T-MGs with > 99% purity by plating the cells on mixed cortical cultures on non-TC-treated plastic and washing with sterile 1X PBS[19, 20] (Supplemental figure 1).

To test whether T-MGs are sufficient to populate the brain parenchyma, we transplanted T-MGs directly into the cortex and hippocampus of untreated wild-type (WT) mice. As expected, and previously demonstrated, injected T-MGs failed to repopulate the mouse brain, as evidenced by the presence of cellular debris carrying GFP signal immediately at the injection site (GFP+/IBA+) (Figure 1A). Furthermore, the injection site was surrounded by a high density of endogenous, highly reactive, microglia cells, which lacked GFP expression but were labelled by IBA1+, microglial marker. The strongest IBA1 signal was maintained up to 1 mm in both directions from the injection site[21] (Figure 1A, B). No GFP+IBA+ cells were observed distally from the injection site (Figure 1A, B). These results further support previous studies, which demonstrated that a robust depletion of endogenous microglial (E-MG) populations is required for a successful replacement of the endogenous microglia to minimize the number of reactive host microglia at the injection site[7, 22].

**Figure 1.**
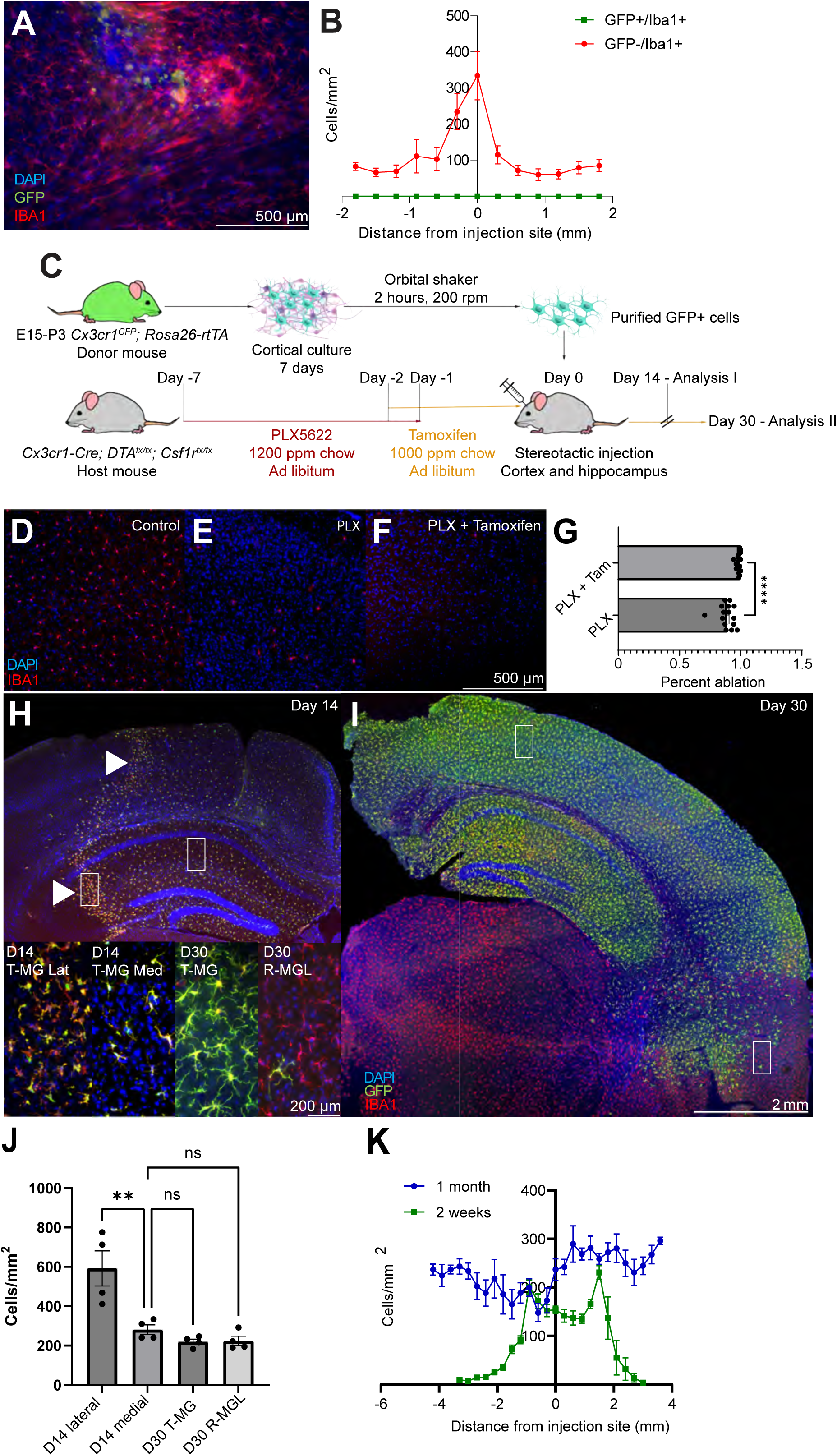
Microglial spread in an ablated host: (A) Representative image of microglia transplanted into an unablated host. (B) Quantification of endogenous IBA1+/GFP-response to GFP+ transplants in an unablated host. (C) Schematic for microglial transplant protocol. (D-F) Images of IBA1+ density within CSF1R cKO host animals at baseline, treated with PLX5622, and PLX5622 + Tamoxifen, respectively. (G) Quantification of IBA1+ cell density set relative to unablated density, n = 10 mice per treatment group. (H, I) Representative coronal sections of a mouse with 200,000 GFP+/IBA1+ cells stereotactically injected into the somatosensory cortex, 2 weeks post-transplantation (H) and 1-month post-transplantation (I). GFP indicates the T-MGs. (J) Quantification of GFP+ cell density both in the center (medial) and within 300 µM of the transplant edge (lateral). (K) Binned density of serial sections in the cortex of the associated post-transplanted mice, measured every 300 µm in sections 300 µm apart. 0 is the site of injection, 1 mm to the right of Bregma. N = 5 mice per treatment group. Statistical significance found via two-way ANOVA with Bonferroni correction, * = p < 0.05, ** = p < 0.01, **** = p < 0.0001.

Next, to confirm that ablation of E-MGs is necessary for T-MGs repopulation, we significantly depleted E-MG using a combination of genetic and pharmacological approaches. For this purpose, we pharmacologically inhibited CSF1R, the signaling receptor critical for microglial growth and survival[23], using PLX5622, a small-molecule CSF1R inhibitor[23], one week prior to surgery. PLX5622 was administered to 4-month-old *Cx3cr1^CreER/+^;Rosa^DTA^;CSF1R^fl/fl^*conditional knockout animals (CSF1R cKOs). This protocol successfully eliminated most microglia, however a few residual ones remained (Figure 1C-E). To ensure complete E-MG ablation and prevent ablation of T-MGs, we administered PLX5622 drug to an adult donor mouse carrying tamoxifen-inducible genetic knockout of CSF1R combined with an overexpression of diphtheria toxin. To maintain E-MGs ablation after PLX5622 withdrawal, 24 hours prior to transplantation, we transitioned the animals to tamoxifen chow. The combined approach further increased microglial ablation to 95% (Figure 1C, F, G).

Using the combined microglia depletion approach, we injected 200,000 cells/µL of GFP+ T-MGs into the cortex and hippocampus of ablated CSF1R cKOs (Figure 1H, I). As previously observed[22], 2 weeks post-transplantation, T-MG (GFP+IBA+) formed a proliferative wave as they retiled the tissue (Figure 1H) that increased microglial density from an average of 271 to 576 cells/mm^2^ (Figure 1J, K). We did not observe the proliferative wave at the one-month post-transplantation time point, when spreading had ceased and the R-MGLs (GFP-IBA+) had repopulated the remaining parenchyma (Figure 1I-K), demonstrating their capacity for efficient repopulation.

### Repopulating microglia differ in phenotype from resting endogenous microglia

As previously reported, cultured microglia have a less ramified morphology[24], and downregulate genes typical to E-MGs, like Tmem119, P2ry12, and Sparc[25]. However, purified microglia in culture have been shown to overexpress markers of peripheral monocytes like Anxa1, along with markers of high proliferation like Birc5 and markers of activated microglia like Lpl[26]. Furthermore, microglia transplanted in postnatal pups are functionally indistinguishable from their in vivo counterparts[27]. Thus, to determine whether our cultured T-MGs regain the profiles of in vivo microglia after transplantation into adult mice, we compared the morphology and expression of CD68, a marker of immune activation[28], between T-MGs, E-MGs, and IBA1+/GFP-host microglial cells (R-MGs) that repopulated the CSF1R cKOs brains (Figure 2). Our results showed that two weeks post-transplantation, only T-MGs located at the periphery of the spreading area had significantly increased CD68 expression, suggesting immune activation (Figure 2A-D).

**Figure 2.**
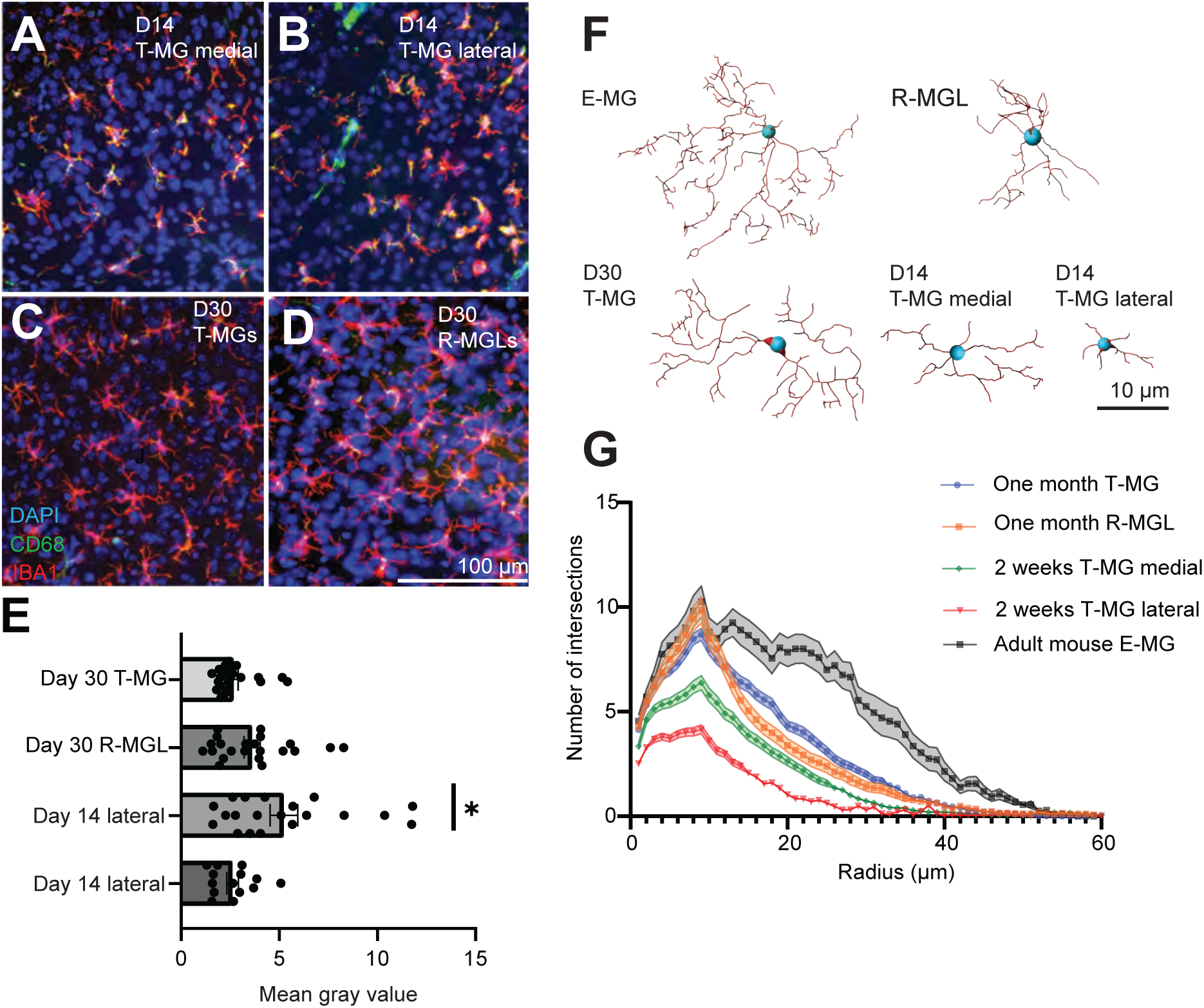
Inflammatory profile of T-MGs, R-MGLs, and E-MGs: (A-D) Representative images of GFP+ cells, both within and without the transplant border (medial and lateral, respectively) 2 weeks pos-transplantation. Cells were compared to the month-old transplant, along with unaffected adult microglia. (E) Quantification of CD68 (cell activation) expression in these cells. (F) Morphological complexity analysis of microglial populations via Sholl analysis, along with 3D wireframe representations of the cells analyzed using Imaris software (K.1-4). (G) Sholl analysis of microglial types. n = 20 cells per mouse (100 cells), n = 5 mice. Statistical significance found via two-way ANOVA, with Bonferroni correction * = p < 0.05.

To assess the branch morphology and complexity of these three microglial populations, we performed 3D reconstruction and Sholl analysis[29] 30 days post-transplantation. The T-MGs morphology continued to show lower ramification than E-MGs, but matched the complexity of R-MGLs (Figure 2F, G), suggesting that repopulating R-MGLs and T-MGs maintain characteristics typically associated with inflammation[30]. Overall, T-MGs repopulated with rates and morphologies similar to R-MGLs, however, both repopulating populations remained structurally distinct from E-MGs (Figure 2F, G).

### T-MGs modified to deliver BDNF retain immune capabilities in vivo

Temporal control of protein payload delivery to the brain is essential in clinical settings to halt the delivery in case of systemic side effects. Previous studies have shown that microglia can deliver a protein payload to the brain[15], but they achieve it without temporal control. Thus, to evaluate our capacity to deliver a temporally regulated protein payload, we engineered microglia to express Brain Derived Neurotrophic Factor (BDNF) throughout the central nervous system through controlled exogenous drug administration.

We chose BDNF as a test payload due to its potential therapeutic benefits in Alzheimer’s Disease (AD) shown in pre-clinical mouse models, where it partially reversed the 5xFAD mouse phenotype when genetically overexpressed in astrocytes[31]. Also, human AD patients with reduced BDNF levels had reduced neuronal densities in the cortex[32]. Notably, at the transcriptional level, endogenous microglia show no detectable levels of BDNF expression[33]. Thus, we constructed an inducible lentiviral vector carrying BDNF under a tetracycline-dependent promoter (Supplemental figure 2A), which, through expression of rtTA at the *Rosa26* locus, allows for temporal control of the *tet* promoter via administration of doxycycline[34].

In vitro, microglia infected overnight with the BDNF-containing lentivirus showed BDNF overexpression in media collected 72 hours after initiating doxycycline treatment (Figure 3A, B). ELISA results indicated a 3-fold increase in BDNF expression in doxycycline-treated conditions compared to control GFP-expressing virus or no virus (Figure 3B).

**Figure 3.**
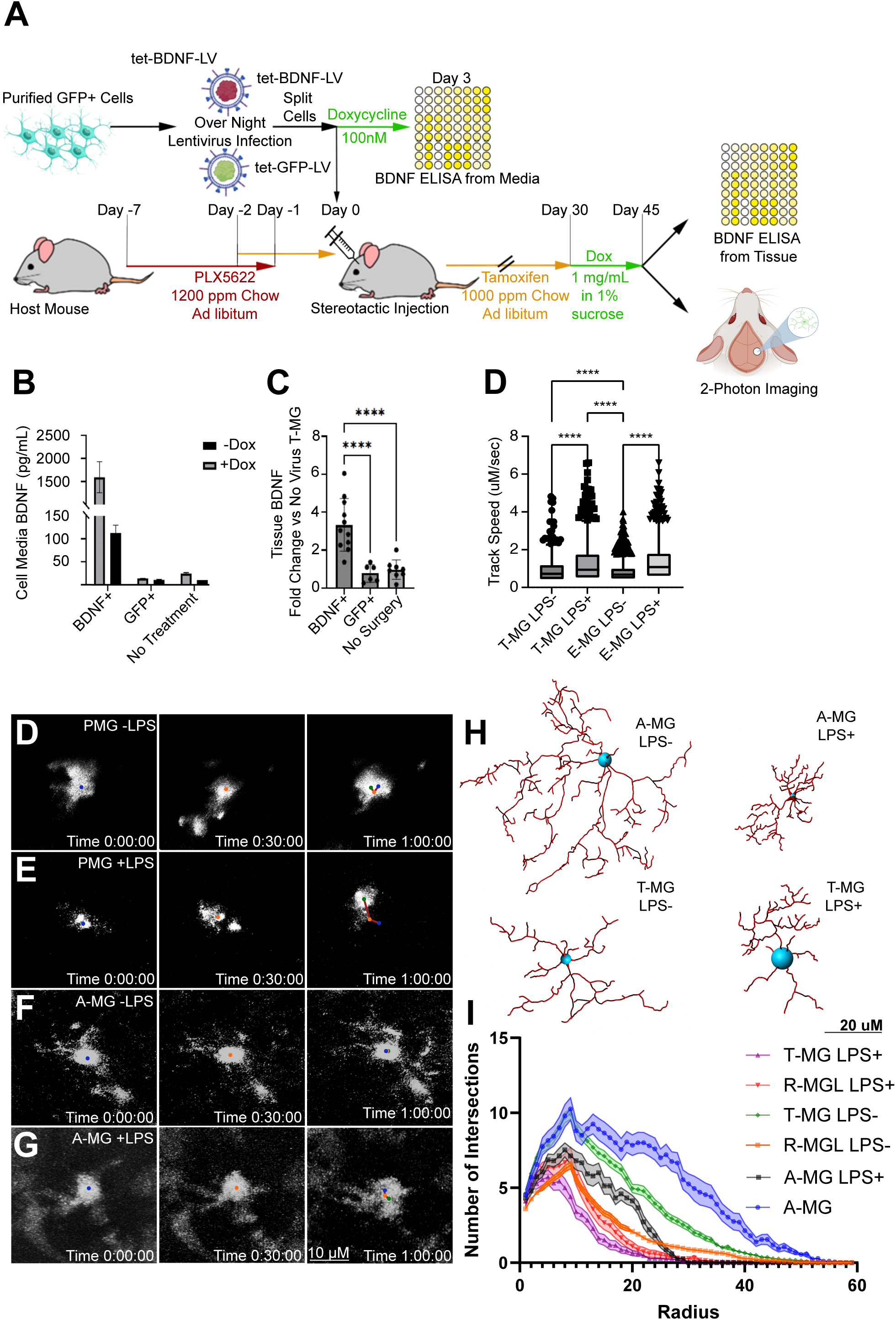
T-MGs retain their functional capabilities in vivo: (A) Schematic for BDNF lentiviral infection of T-MGs, along with illustration of downstream analysis. (B) ELISA run on media harvested from T-MGs in vitro following 72 hours of doxycycline exposure. (C) ELISA run on tissue lysates harvested from ideal host animals one month after surgery, with the animals being put on doxycycline water for one week. n = 8-10 mice per treatment group. D) Quantification of track speed from two-photon images. (E-H) Representative frames of two-photon imaging, along with track lengths highlighted. (I) Representative immunohistochemistry images from transplanted and control cells processed via Imaris. (J) Quantification of cellular complexity analyzed via Imaris Sholl Analysis plugin, n = 5 mice per treatment group, 50 cells analyzed per mouse. Analysis performed via two-way ANOVA with Bonferroni correction. **** = p < 0.0001

To test our approach in vivo, we ablated endogenous microglia in CSF1R cKOs and transplanted T-MGs transduced with the tetracycline-dependent BDNF lentivirus (tet-BDNF-LV) into the mouse cortex and hippocampus. To induce BDNF expression, the animals were maintained on tamoxifen for 30 days to ablate endogenous microglia, followed by two weeks of doxycycline-treated water to induce BDNF expression in T-MGs (Figure 3A). IHC analysis of transplanted animals revealed that T-MGs were present throughout the cortex (Supplemental figure 2B-E).

To assess BDNF expression levels, we collected tissue biopsy punches 0.5 mm posterior to the injection site and performed ELISA (See Methods). Infected T-MGs showed a threefold increase in BDNF when doxycycline was administered, as compared to non-infected T-MGs (Figure 3C), suggesting a robust induction and expression of the payload protein.

Next, we examined whether transplanted tet-BDNF-LV-infected microglia could respond to pro-inflammatory stimuli. One month after transplantation, we stimulated microglia with LPS and imaged their responses in vivo, using 2-photon microscopy (Figure 3A). We observed that 24 hours post-LPS stimulation (1 mg/kg IP), E-MGs within WT control mice showed higher motility compared to those same cells prior to LPS stimulation (Figure 3 D-F). However, T-MGs were more motile compared to E-MGs even at rest (Figure 3D, G, H), and exhibited greater loss of ramification than E-MGs following LPS stimulation (Figure 3I, J).

These findings suggest that in vivo, T-MGs retain reactivity to damage, and lentiviral modification does not impede their inflammatory response when delivering a drug in the CNS. However, baseline motility differences did appear to exist, aligning with the morphological divergences observed on immunohistochemistry (Figure 2G).

### Transcriptomic analysis of microglial transplants infected with BDNF lentivirus

To further ensure T-MGs specifically expressed BDNF and that lentiviral expression does not significantly alter gene expression profiles, we performed single-cell RNA-sequencing on three sample groups: *CSF1R* cKOs transplanted with microglia with or without BDNF lentivirus (BDNF or NoV), and age-matched *Cx3cr1^GFP^;Rosa26-rtTA* adult mice (WT) without T-MGs transplantation, two months post-transplantation of T-MGs into ablated *CSF1R* cKOs. We sagittally sliced the whole brain samples and dissociated the left hemisphere into single cells for RNA sequencing, while the right hemisphere was used for cell engraftment verification.

Quantification of brains with transplanted microglia with or without the BDNF virus contained both T-MGs and R-MGLs (Supplemental figure 2D), with tissue consisting of approximately 60:40 ratio of GFP+/IBA1+ to GFP-/IBA1+ cells (Supplemental figure 2E). Similar to Figure 2F, both T-MGs and R-MGLs were morphologically similar (Supplemental figure 2F). All cell types from whole-brain samples were initially subclustered using the ScType Classification algorithm[35] to identify microglia (Supplemental figure 3A, B), which were then computationally isolated and subclustered for further analysis. Cell clusters were grouped separately by origin from transplanted or non-transplanted brains (Figure 4A). We observed that T-MGs/R-MGLs transplanted in CSF1R cKO host mice clustered together, but separately from E-MGs from donor WT mice that did not receive transplanted microglia (Figure 4B), suggesting common molecular mechanisms used by T-MGs and R-MGLs.

**Figure 4.**
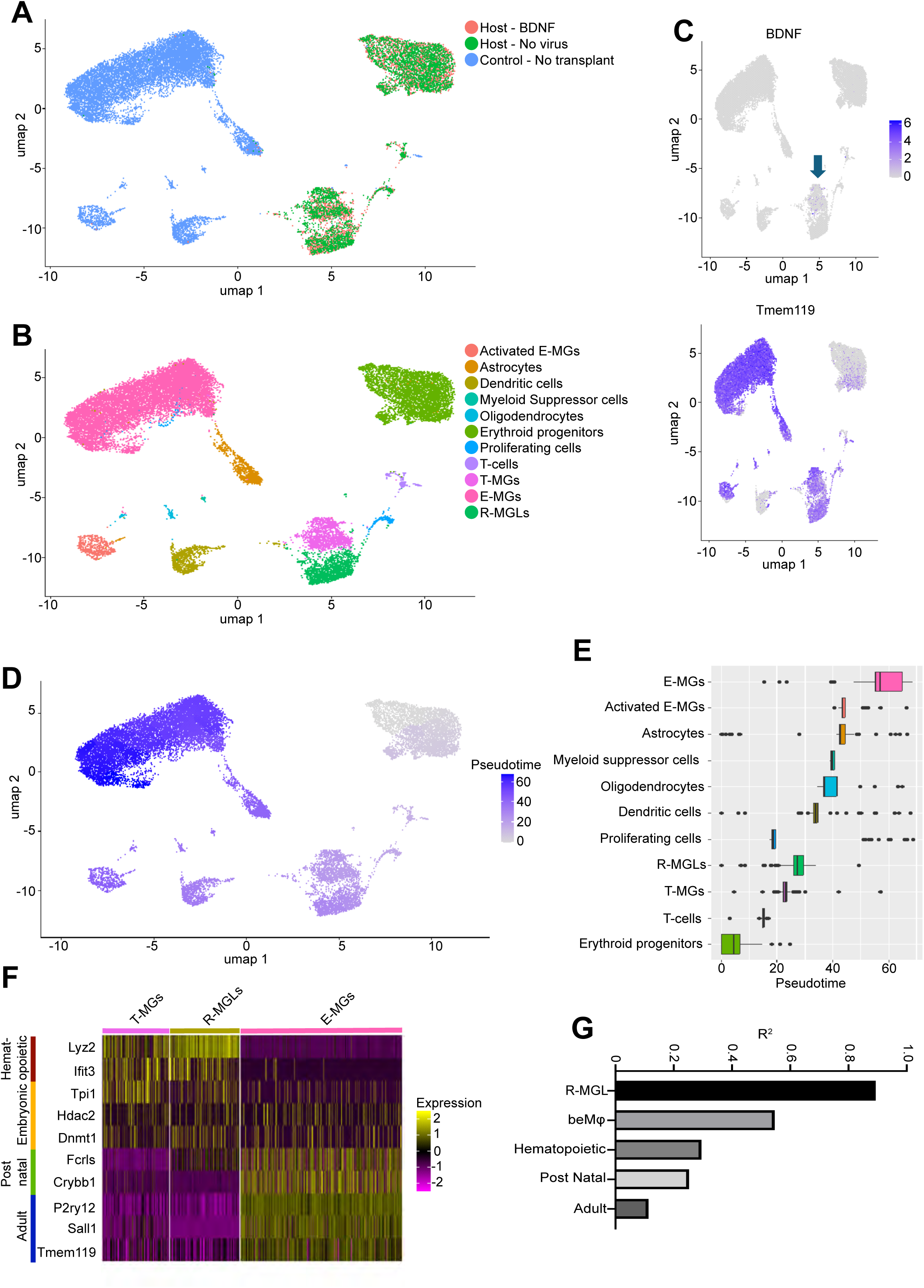
Transcriptional analysis of T-MGs: (A) UMAP Projections of microglial-like cells identified by the sctype sorting algorithm, split into three treatment groups: unaffected age-matched adult *Cx3cr1^GFP^;Rosa26-rtTA* mice, *CSF1R* cKO host mice transplanted with T-MGs that were infected with the BDNF lentivirus (BDNF), and host mice transplanted with T-MGs without the virus. (B) UMAP Projection broken down by cell group. Markers for these categories can be found in Supplemental Figure 3B. (C) Feature plots of BDNF, which is localized to one specific cluster highlighted by the arrow, and TMEM119, a marker for microglia. (D, E) Pseudotime analysis via monocle3, broken down as Feature Plot and a graphical representation of scores broken down by cluster designation from (B). (F) Heatmap analysis, binned by T-MG, R-MGL, or E-MG. T-MGs and R-MGLs showed much higher incidents of embryonic microglial markers (Hdac2, Tpi1), while E-MGs express markers of adult resting microglia (P2ry12, Tmem119). (G) Comparison of R^2^ scores of T-MG vs Postnatal, Embryonic, and Adult Microglial genetic signatures, along with beMφs[40]. n = 4 control and 4 transplanted animals. Transplants were split into two without the virus and two BDNF-infected T-MG groups.

To further confirm that our T-MGs infected with BDNF lentivirus express higher levels of BDNF, we represented cells from the three hosts into TMEM119 (a microglia-specific marker[36]) positive and TMEM119 negative clusters. We observed three main TMEM119-positive clusters, one containing E-MGs and erythroid progenitors, the second containing activated E-MGs, and the third containing T-MGs and R-MGLs (Figure 4C). Significantly, only the third cluster contained T-MGs and R-MGLs, which showed significant BDNF overexpression, specifically in the T-MG population (Figure 4B, C), further supporting that our approach induced BDNF expression in T-MGs at high levels.

Pseudo-time analysis placed T-MGs and R-MGLs on similar pseudotime points (Figure 4D, E), suggesting molecular similarities. These two clusters were enriched for genes such as *ApoE*, *Lyz2*, *Ms4a7*, *Clec12a*, *Ifitm3*, and *Igf1* (Figure 4F), genes previously shown to be upregulated in repopulating microglial-like cells, embryonic microglia, and brain border macrophages[37, 38]. T-MGs and R-MGLs also expressed genes more specific to embryonic microglia sampled from E13-E15 cortical tissue, such as *Tpi1*, *Hdac2*, and *Dnmt1*[37] (Figure 4F), suggesting likely not-yet completed maturation during brain repopulation. As expected, E-MGs expressed genes more typical of adult microglia: *TMEM119*, *P2Ry12*, *Sall1*, and *Fcrls*[39] (Figure 4F).

Differential gene expression in T-MG and R-MGL clusters was highly correlated relative to WT E-MGs (Supplemental figure 3E), indicating that recently repopulated microglia, whether T-MGs or R-MGLs, closely resemble each other. Within T-MG clusters, there was a strong correlation between cells with and without BDNF expression (Supplemental Figure 3F), suggesting that viral modification does not affect the transcriptomic profile. No sex differences were observed (Supplemental figure 3G).

Comparison with published embryonic, postnatal[37], and bone marrow-derived macrophage[40] datasets revealed that T-MGs and R-MGLs correlate closely with bone marrow-derived macrophages (beMφs) (Figure 4G, Supplemental figure H). Our results indicate that T-MGs, though transcriptionally distinct from E-MGs, possess a similar transcriptome profile to other R-MGLs, which repopulate the brain following ablation. In chronic neuroinflammation, these R-MGLs may benefit the CNS[41], adding to the rationale for exploring T-MG therapeutic applications in adult wild-type mice.

### Multiple rounds of host microglia ablation with PLX5622 allow T-MGs in WT mice to spread

To mimic the treatment structure used in a clinical setting, we transplanted BDNF-infected microglia into the parenchyma of WT animals that underwent only pharmacological microglial ablation prior to transplantation. As shown in Figure 1A-B and in previous studies[14, 15], establishing a physiological niche for microglial replacement requires the ablation of host microglia. In addition, residual host cells that survive depletion tend to proliferate in response to damage around the injection site of the transplanted microglia, inhibiting their survival or spread (Figure 5C). Although genetic ablation of host microglia in the method described above provides a niche for recolonization of the tissue by transplanted microglia, for clinical relevance, it is crucial to transition away from hosts with genetic microglial ablation to hosts with pharmacological ablation. Previous methods without genetic ablation have used lethal dose irradiation or chemotherapy of the host to allow replacement of host microglia with transplanted microglia[7, 12, 13].

**Figure 5.**
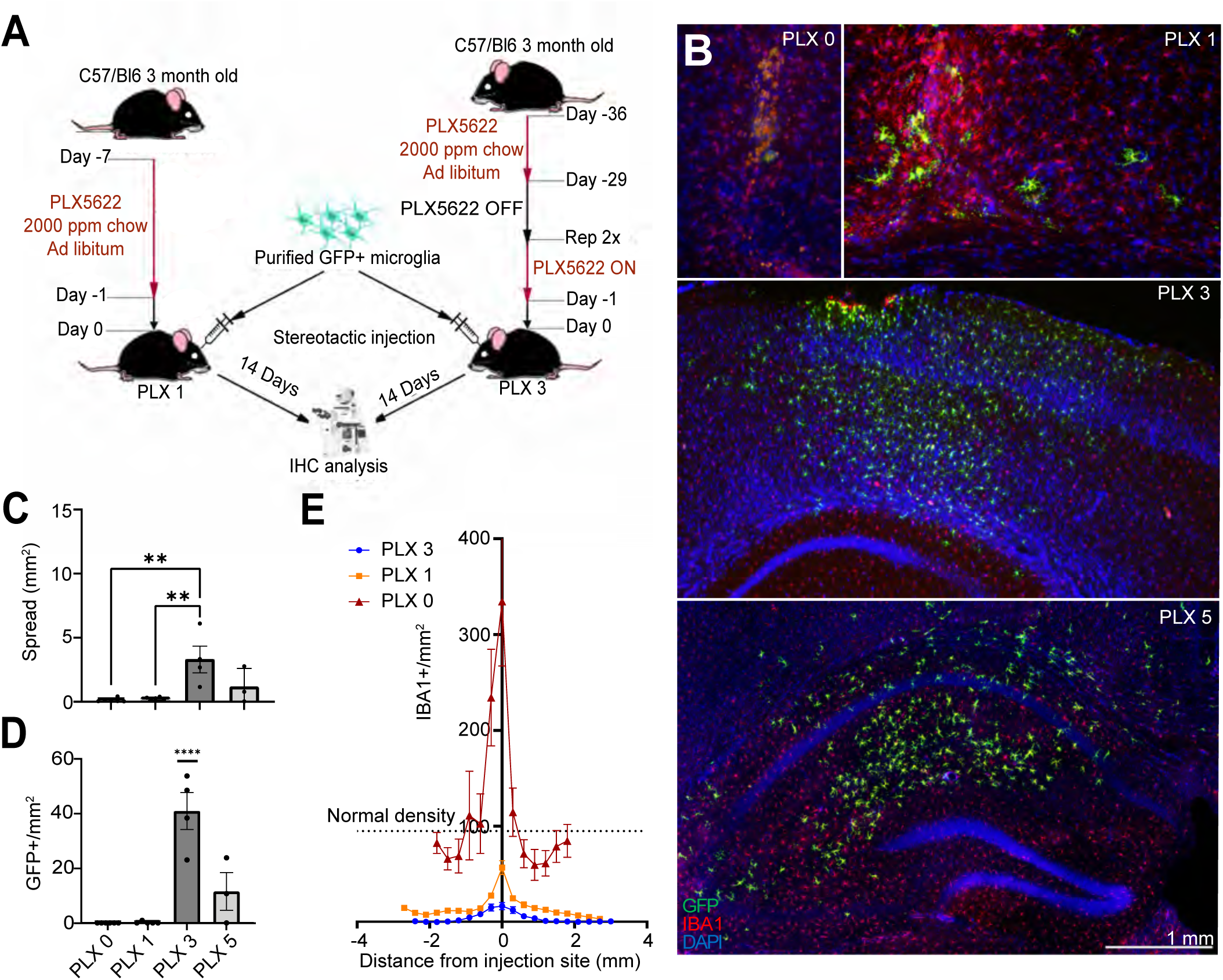
Pulsatile PLX5622 serves to facilitate T-MG spread in WT animals: (A) Schematic for microglial transplant protocol and pulsatile PLX5622 dosing regimen. PLX3 represents 3 rounds of PLX5622 dosing at 2000 ppm, one week on followed by one week off. (B) Representative images from microglial transplants into 3-month-old WT mouse subject to various numbers of PLX5622 pulses, each transplant contained 200,000 cells, along with quantification of the area spread. (C) Density of IBA1+, GFP-cells following an injection of cells with increasing numbers of PLX5622 pulses. (D) GFP+ cell density within the transplanted area from each PLX5622 dosing regimen. (E) GFP+ area spread quantification in each of the pulsatile experiments. n = 3 animals for PLX56225 treatment, n = 5 for all other treatment groups. Analysis via two-way ANOVA with Bonferroni correction. * = p < 0.05.

We found that a one-week course of CSFr1 inhibition with PLX5622 at 2000 ppm before transplantation was sufficient for replacement when combined with the genetic ablation; however, it was insufficient for cell engraftment in WT animals (Figure 1A, PLX1). This is due to the rapid repopulation by endogenous cells following the cessation of PLX5622 (necessary to avoid also killing transplanted microglia), which prevents the spread of transplanted cells (Figure 5B, 2C). Thus, we targeted resident progenitors with multiple rounds of PLX5622 administration, similar to PLX 3397 regimen[14]. We found that WT animals dosed with intermittent PLX5622 pulses every other week for three (PLX 3) to five cycles (PLX 5) prior to transplantation (Figure 5A) showed a substantial increase in the grafted cell area and cell density within the grafted area (Figure 5C, D). This was paired with a significant decrease in R-MGL cell density (IBA+ cells/mm^2^) around the injection site (Figure 5E). Taken together, these findings suggest that providing three pulsatile doses of PLX5622 prior to transplantation on adult WT animals efficiently eliminates residual E-MGs and R-MGLs, which further limits their response to stereotactic injection damage, enabling successful microglia transplantation and wider spread from the injection site in these hosts.

### A modified constitutively active CSF1R increases the rate of T-MG spread

While the triple iterative PLX5622 treatment protocol (PLX 3) enabled successful engraftment of transplants in WT animals, the area of spread was significantly limited compared to transplants in genetically modified hosts (Figures 1C, 4B). With the speed at which microglia repopulate the CNS even in the setting of triple PLX5622, we aimed to design a virus that could confer a competitive edge to the T-MGs during repopulation in a doxycycline-dependent manner. We hypothesized that this approach would allow faster T-MG repopulation due to limited R-MGL competition.

To enhance the spread of and give the T-MGs a survival advantage, we designed a lentivirus with a mutated *CSF1R* (Supplemental figure 4A), incorporating mutations L301S and del706-712. The L301S mutation allows spontaneous receptor activity[42], and del706-712 prevents receptor internalization post-activation[43]. This mutant receptor was placed under a tetracycline-specific promoter, enabling temporal expression of CSF1R only in the presence of doxycycline.

In vitro, engineered cells transduced with mutated *CSF1R* reached higher densities after two weeks on mixed cortical cultures compared to those transduced with a doxycycline-dependent mCherry control virus (Supplemental figure 4B-D), suggesting a competitive advantage of transduced microglia. Due to the size of the *caCSF1R* construct, no fluorescent tag was attached to the caCSF1R virus, due to limited lentivirus capsid capacity. However, we designed the mCherry control virus with the same viral backbone (Supplemental figure 4B) and infected cells in the same fashion to use as an approximation for construct expression. In *CSF1R* cKO mice without doxycycline treatment, we observed 5% of infected T-MGs expressing mCherry, while in mice with doxycycline, the T-MG mCherry expression rate climbed to 90% (Supplemental figure 4E - G). Thus, we transplanted *CSF1R* cKO host mice with 50,000 cells transduced overnight with *caCSF1R* or *mCherry* control virus. Brains were collected and analyzed three weeks later (Figure 6A).

**Figure 6.**
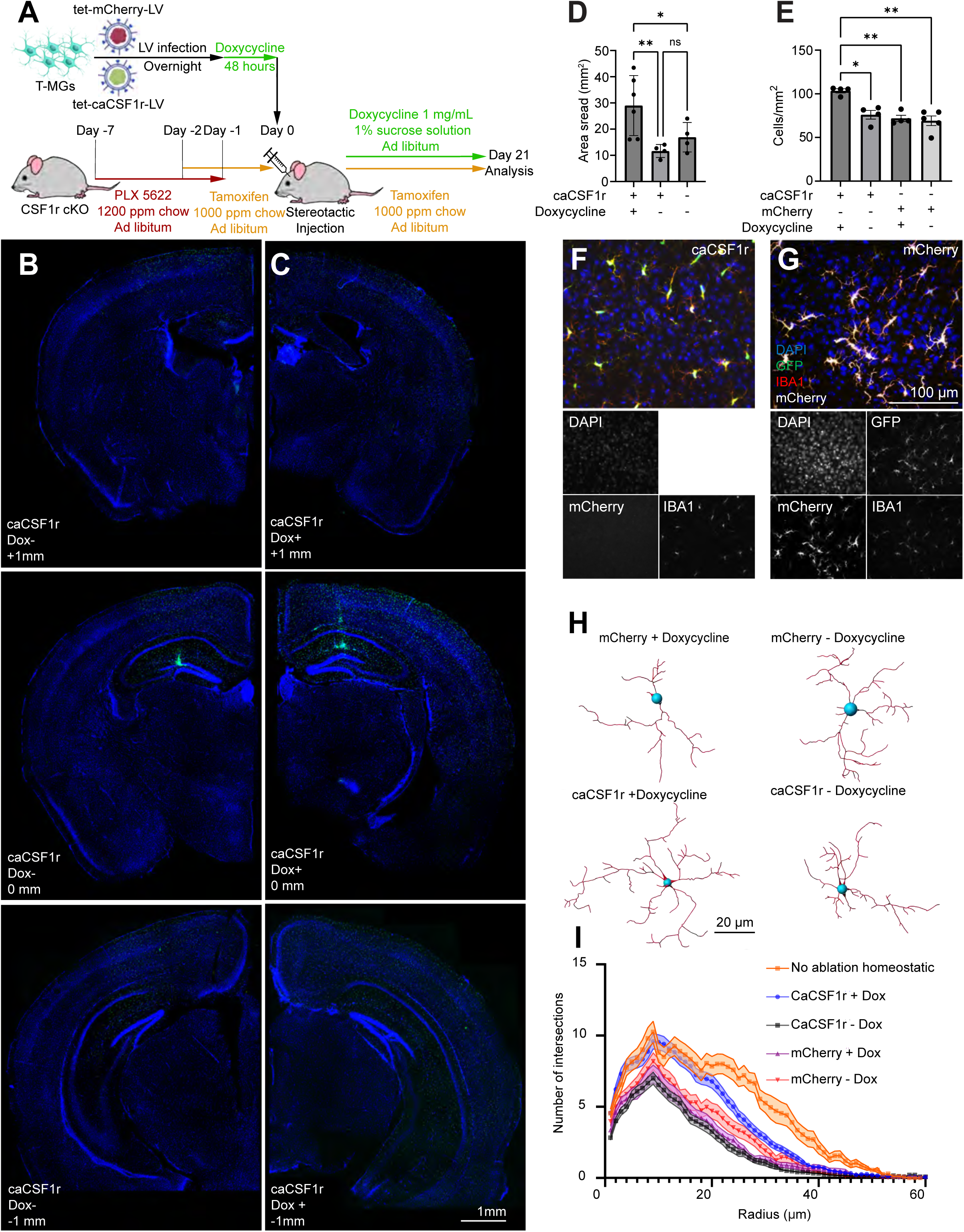
caCSF1R construct confers increased spread in a doxycycline dependent manner: (A) Schematic for microglia transplant with caCSF1R virus. (B, C) Representative images of transplanted cells infected with *caCSF1R* lentiviruses, transplanted in animals treated with or without doxycycline infused water, 3 weeks post-transplantation. Images were taken at the injection site, along with 1mm anterior and 1mm posterior. (D) spread quantification of transplanted cells following three weeks in vivo compared to transplanted uninfected cells following one month in vivo. (E, F, G) Quantifications of cellular density within cell transplants, along with representative images of caCSF1R (F) and mCherry (G) transplants in doxycycline administered animals . N = 4 for all treatment groups. Analysis via two-way ANOVA. (H) Representative images of Imaris 3D reconstructions between caCSF1R infected and mCherry infected T-MGs with and without doxycycline treatment. (I, J) Sholl quantification of cellular complexity per treatment, with Sholl sums compared in (I) and Sholl intersections per intersection illustrated in (J). N = 10-20 cells per mouse, N=4-5 mice per treatment. Statistical analysis: Kruskal-Wallis test and Dunn’s test for multiple comparisons.

Three weeks post-transplantation, T-MGs transduced with mutated *CSF1R* spread on average 300% further from the injection site in mice treated with doxycycline than those that were not treated with doxycycline (Figure 6B-D). This spread is not only doxycycline dependent, but significantly improves upon the engraftment rates observed in uninfected transplants (Figure 6D). This increased spread was accompanied by a higher microglia density within the area of spread (Figure 6E-G). Notably, the complexity of microglial morphology appeared to increase in a doxycycline-dependent manner in T-MGs transduced with the *caCSF1R* virus as compared to controls (Figure 6F-J). The use of caCS1R virus, although viable in promoting an increase in spread when induced via doxycycline, did not appear to confer any resistance to PLX when compared to mCherry controls (Supplemental Figure 4H, I). This led us to conclude that the *caCSF1R* virus could serve as an additional bolster to microglia spreading and ramification in a WT host.

### T-MGs modified with caCSF1R can deliver BDNF to the brain parenchyma of adult WT mice

After establishing a robust PLX5622 treatment regimen and endowing T-MGs with a doxycycline-inducible competitive edge over endogenous cells, we hypothesized that these microglia could provide effective BDNF delivery to the parenchyma of WT animals that was applicable to the clinic. To test this possibility, we infected microglia with *caCSF1R* virus and delivered them via stereotactic injection into the somatosensory cortex of 3-month-old WT animals pretreated with three one-week pulses of PLX5622 (Figure 7A). When comparing microglial spread in WT animals, those that were infected with *caCSF1R* spread approximately threefold further than those without the *caCSF1R* virus (Figure 7B, C). Additionally, these cells appeared to exhibit higher ramification when compared to non-*caCSF1R* infected T-MGs as quantified via Sholl analysis (Figure 7D, E). Following these analyses, we transplanted microglia infected with both BDNF and caCSF1R lentiviruses to examine whether we could deliver functional BDNF to the parenchyma of WT animals pretreated with three pulses of PLX5622.

**Figure 7.**
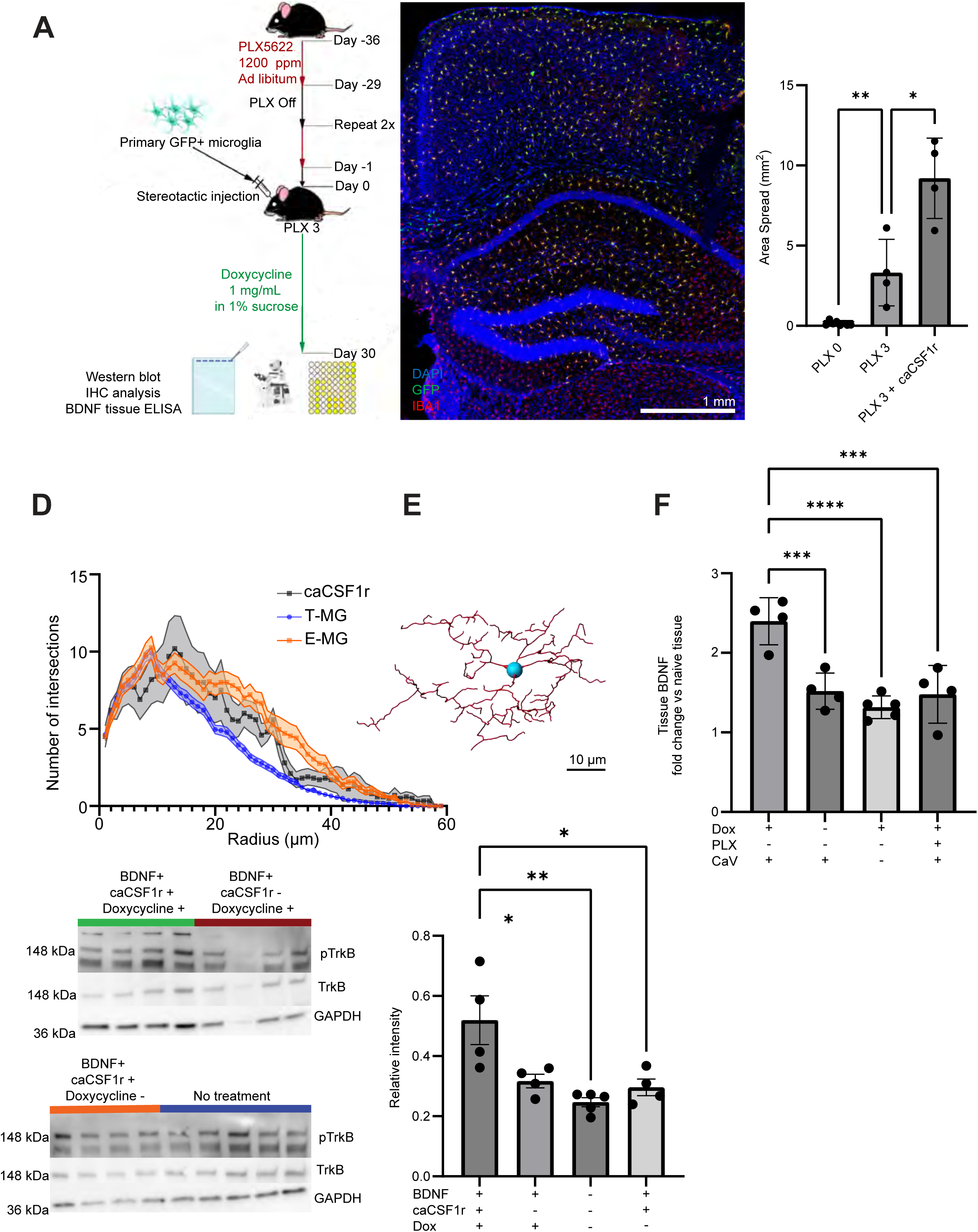
*BDNF/caCSF1R* infected T-MGs can deliver BDNF to the parenchyma of WT animals in a doxycycline dependent manner: (A) Schematic for microglial transplant protocol using the combination of pulsatile PLX5622 and *caCSF1R* virus. (B) Representative image of microglia transplanted with *caCSF1R* virus transplanted within WT cortices undergoing the described treatment regiment. (C) Quantification of spread in WT animals not pretreated with PLX, treated with 3 pulses of PLX, and treated with 3 pulses of PLX with microglia infected with *caCSF1R* virus. (D) Sholl analysis between transplanted cells in the conditional knockout host compared to transplant in the B6 mice. (E) a wire diagram representation of a transplanted *caCSF1R+* cell. (F) ELISA data of tissue biopsy punches from B6 mice transplanted with microglia infected with both caCSF1R and BDNF viruses, with and without doxycycline. n = 4 for all treatment groups. (G) Image of Western blots, assessing levels of phosphorylated Trk B, total TrkA, total TrkB, and GAPDH. (H) Quantification of pTrkB expression, as fold change over TrkB expression within the sample. Analysis via two-way ANOVA with Bonferroni correction, n = 4-5 mice per treatment group.

ELISA analysis of tissue biopsy punches taken from these WT mice 0.5 mm from the transplant revealed increased BDNF levels in hosts with T-MGs infected with both *caCSF1R* and *BDNF* viruses in WT mice treated with doxycycline than in control animals (Figure 7F). Both doxycycline and the *caCSF1R* virus were necessary for BDNF delivery. Importantly, *caCSF1R* negative microglia did not alter baseline levels of BDNF, and neither did microglial transplants without doxycycline administration (Figure 7F). Western blot analysis revealed that WT animals transplanted with microglia co-infected with the *caCSF1R* and *BDNF* viruses exhibited an increased level of pTrkB (a marker of BDNF activity[44]) in a doxycycline controllable manner (Figure 7G, H). Compared to the control group, mice with the transplant had a two-fold increase in the intensity of the pTrkB band, demonstrating a functional change in the host brain brought about by our engineered microglia.

Overall, our results suggest that the microglia cell transplantation strategy described here could, in principle, be used to deliver inducible levels of therapeutic biologics throughout the brain in a clinically relevant manner.

## DISCUSSION

Our study developed a methodology to engineer primary T-MGs as delivery vehicles for temporally controlled drug delivery into the CNS. We demonstrated that our microglia replacement method can serve as a sustainable approach for widespread CNS delivery of brain-derived neurotrophic factor (BDNF). T-MG showed elevated BDNF expression in the brain parenchyma of the host and increased pTrkB expression, demonstrating the delivery of a biologically active product. In addition, we found that combining a pulsatile PLX5622 dosing of the host with tetracycline-dependent expression of a constitutively active CSF1R in the donor microglia allows the introduction of genetically engineered microglia into a host immune environment without the need for aggressive conditioning. These findings further advance the goal of making microglial transplants a more clinically feasible methodology for CNS drug delivery.

We also identified key differences between transplanted microglia and homeostatic endogenous microglia, while highlighting the similarity of T-MGs to repopulating microglia-like cells within the CNS. Our approach provides a viable model for drug delivery to the brain in wild-type animals, suggesting that microglial replacement could significantly advance neurological treatments. The ability to genetically modify microglia offers the potential to deliver therapeutic proteins effectively throughout the brain. Previous studies have indicated that microglia subjected to in vitro culture conditions exhibit distinct gene expression profiles compared to their in vivo counterparts. This is the case both for primary[45] and hiPSC derived cells[46].

Xenotransplanted microglia placed in the brain of early postnatal pups appear to align more closely with their ex vivo counterparts[47]. However, studies that replace microglia in adult animals mainly focus on the morphology of the engrafted microglia[48], or use animals lacking endogenous microglia, so there are no R-MGLs to compare[22]. Transcriptome data obtained in this study from transplanted and non-transplanted hosts further confirmed the feasibility of the approach to express and deliver BDNF throughout the brain while maintaining temporal control on the expression.

In our analysis, T-MGs were found to retain distinct morphological and transcriptomic profiles, more closely resembling R-MGLs than E-MGs. Specifically, T-MGs showed lower levels of homeostatic microglia markers such as Tmem119, P2ry12, and Sall1, while displaying elevated levels of embryonic and hematopoietic markers, including Lyz2 and Tpi1. These findings align with previous replacement studies and emphasize the potential of exogenous primary microglial cultures [22],[49]. Despite these differences, T-MGs still responded to pro-inflammatory stimuli and ceased proliferation upon encountering IBA1+ cells, showcasing their therapeutic viability. Furthermore, our findings show that lentiviral transduction with the tet-BDNF construct did not alter the transcriptomic or morphological characteristics of T-MGs. This stability enabled us to further investigate T-MGs functional potential to deliver BDNF in healthy adult animals.

Our study did not track lineage origins, which limited our ability to determine if R-MGLs derived from resident microglia or bone marrow. Comparisons to previous studies[40] revealed similarities between R-MGLs, T-MGs, and blood-derived macrophages. Future research should consider different orally available drugs, like rapamycin[50], to separately control constructs for transplant spread and drug delivery. Longer-term studies, extending beyond the current 2-month scRNA timeframe are essential to fully assess the sustainable impacts of this delivery technique. Additionally, long-term viability studies of caCSF1R infected microglia should be undertaken, as initial findings suggest that they take on a morphology more similar to E-MGs when compared to their noninfected counterparts. This would also allow for further investigation into whether the T-MGs and R-MGLs converge with E-MGs.

Some studies investigating microglial transplants rely on extensive immune modulation through immunodeficient animals[51] or by employing full-body irradiation/chemotherapy drugs[7]. A recent study reported that microglial replacement can deliver the protein neprilysin to the brain of transgenic AD mice with the immunocompromised MITRG background[15]. Our study advances on these methods by demonstrating a doxycycline-dependent delivery of a protein to WT mice, instead of immune-compromised mice. The transition to a minimally invasive regimen of small-molecule pharmacotherapy before transplantation is crucial to maximize the clinical applicability of microglial replacement. PLX5622 is highly selective for CSF1R, sparing similar immunomodulatory targets[52]. This allows for the transient removal of microglia without long-lasting effects to the remainder of the immune system, as evidenced from a previous study[53] and our observations that even after multiple rounds of PLX5622, a new microglial-like population recovers. The continuous ablation strategy of tamoxifen may play a role in the deviations observed between T-MGs and R-MGLs, as a new study demonstrated a much higher degree of similarity in primary microglia transplanted after three rounds of PLX3397[14].

This study <u>als</u>o builds on the previous publications by administering a biologically active compound using the triple PLX5622 approach. Utilizing T-MGs with an active CSF1R receptor linked to a tetracycline promoter facilitated their integration into the immune-intact adult CNS environment, a considerable challenge for cell transplants. Enhancing the spreading capacity of microglia with *caCSF1R* holds significant therapeutic potential for improving patient outcomes. This approach facilitates more rapid microglial replacement than uninfected microglia or microglia without doxycycline-driven induction of caCSF1R expression. The enhanced spreading speed provides an advantage to T-MGs as they compete with R-MGLs for the repopulation of the host brain. Additionally, the faster spread could serve to reduce the duration for which patients need to maintain immunosuppressive therapies, ultimately minimizing associated risks [54]. Additionally, T-MGs appeared to benefit from CSF1R overexpression and transient microglial ablation, as these cells appeared more highly ramified in both CSF1R cKOs and WT animals. This suggests that CSF1R could serve a similar role in adults as it does during development in differentiating microglia within the CNS[55]. Nevertheless, spread in WT animals was more limited when compared to CSF1R cKO host animals, although T-MGs maintained the capability to deliver BDNF in a doxycycline-dependent fashion. In summary, our study establishes a translationally relevant platform for microglia-mediated delivery of therapeutic biologics, enabling widespread distribution of T-MGs and controlled delivery of a neurotrophic factor in the adult CNS of immunocompetent hosts. These findings highlight the potential of microglia as customizable delivery platforms for neurological therapies. Future work should explore long-term safety to further advance microglia-based interventions for neuroinflammatory and neurodegenerative diseases.

## MATERIALS AND METHODS

### Mouse husbandry

All procedures were approved by the Institutional Animal Care and Use Committee (IACUC) of the Albert Einstein College of Medicine (Protocol #:00001197). Experimental animals were housed in a non-barrier facility in cages maintained at 23 °C. Animals were provided with food and water ad libitum. All rooms were maintained on a 12-hour light/dark cycle, and no more than 5 animals were co-housed except for nursing dams. Cages were changed every week and animals were regularly monitored.

*CSF1R^fl/fl^* (Jax Strain #021212), *Rosa-DTA* (Jax Strain # 009669), and *Cx3cr1^CreER^* (Jax Strain # 020940) animals were crossed to create the *Cx3cr1^CreER/+^;Rosa^TA^;CSF1R^fl/fl^* host animals used for the initial microglial transplants. Primary mouse microglia were derived from *Cx3cr1^GFP^* animals (Jax Strain #005582) crossed with *Rosa26-rtTA* mice (Jax Strain #005670) to generate *Cx3cr1^GFP^;Rosa26-rtTA* microglia for doxycycline inducibility. All other animals were WT C57/Bl6 animals (Charles River Strain #027). Due to the documented sex differences in response to tamoxifen[56], only male WT animals were used. All other hosts were age and sex matched.

### Pharmaceutical compounds

PLX5622 (Chemgood) was used at either 1,200 or 2,000 ppm as indicated in the text. Tamoxifen (Medchem Express HY-13757A) was used at 1,000 ppm. Both drugs were suspended in AIN 76-A powdered chow (Research Diets, D10001); control animals were given AIN 76-A powdered chow without drug during the experimental period. Doxycyline (VWR, 76345-788) was suspended in 1% sucrose (Sigma, S0389) water at 1 mg/mL. All food and water treatments were provided ad libitum. The same drugs were used for in vitro studies at the concentrations noted.*Cx3cr1^CreER^;Rosa-DTA;CSF1R^fl/fl^*animal specific treatment schedule.

In order to ablate endogenous microglia, mice were pretreated with PLX5622 mixed at 1,200 ppm in AIN-76A chow prior to the surgery date. 48 hours before surgery, tamoxifen was added to the chow. 24 hours prior, PLX5622 was removed from the chow. Mice were maintained on tamoxifen for the length of the experimental trial period, as indicated in the text. Doxycycline was administered one month following surgery for tet-BDNF lentiviral induction.

### WT animal specific treatment schedule

Mice were pretreated with PLX5622 at 2000 ppm for varying amounts of time as described in the text. In all cases, PLX5622 was removed 24 hours prior to surgery. Mice were started on doxycycline 48 hours prior to transplantation, and were maintained on doxycycline throughout the trial period to induce activation of the two tetracycline-dependent lentiviral constructs: BDNF and caCSF1R.

### Microglia cell culture

Embryonic day 15 to postnatal day 3 (E15-P3) *Cx3cr1^GFP^;Rosa26-rtTA* cortical tissue was dissociated with MACS Neural Tissue Dissociation Kit (Miltenyi, 130-092-628) according to the manufacturer’s instructions Microglia were isolated using the following adapted protocol[40]: Mixed cortical cells were plated in T75 tissue culture flasks pre-coated with Poly-D-Lysine (Fisher, A3890401) at 0.1 mg/mL suspended in UltraPure Distilled Water (Fisher 10977015) and maintained in microglia enrichment media (Sciencell, 1901) for one week. During this time, the media was changed once after 24 hours to remove debris. Flasks were then agitated via orbital rotation at 200 rpm for 2 hours, and the media containing detached cells was collected and plated on non-TC treated plastic in the same media. One wash was performed one hour following plating, then the media was changed every 3-4 days. Microglia were used for transplantation within one week to avoid cellular senescence. All cultures were maintained at 37 °C with 5% CO2.

### Viral infection

Custom lentiviruses containing the BDNF, caCSF1R, mCherry, and GFP constructs (Supplemental figures 2, 4) were generated by VectorBuilder. Isolated microglia were treated with the appropriate tittered dose of virus added to the media for 16 hours before being collected for transplantation. For microglia transduced with the caCSF1R virus, doxycycline was administered at 100 nM in cell culture media for 48 hours following viral infection prior to transplantation.

### Microglia transplantation

Microglia were detached using Accutase (Corning, 25-058-CI) for twenty minutes, then centrifuged at 300 rpm for 5 minutes. Cells were resuspended at 200k cells/µL in base microglia media without supplements or serum. Mice were placed under continuous anesthesia using isoflurane and secured to a head frame. Cell suspension was placed in 5 µL non-beveled 30 gauge Hamilton syringe (87919) and transplanted in either the hippocampus (+/-1.2 mm, -1 mm, -1.8 mm from Bregma) or the cortex (+/- 1.2 mm, -1 mm, -0.8 mm from Bregma) as outlined in the text. The needle was cleaned with two washes of sterile saline followed by two washes of 70% ethanol, then two washes of sterile saline between animals. For two-photon experiments, microglia were placed more anterior (+/- 1.2 mm, 0.5 mm, -0.8 mm) to avoid interfering with coverslip placement. The cell suspension was slowly injected at a rate of 0.2 µL/min over 10 minutes. Mice recovered on heating pads and were given 5 mg/kg of Meloxicam (Metacam, Boehringer Ingelheim) subcutaneously following the surgery for analgesic purposes.

### Immunohistochemistry

Animals were perfused, as specified by the analysis time point across all experiments, first with cold PBS, then 4% paraformaldehyde (PFA; wt/vol) in PBS. Tissue was post fixed for 16 hours in 4% PFA at 4 °C prior to sectioning using a Vibratome (Leica VT1200S). Sections were 30 µm thick and mounted on Superfrost Plus Slides (Fisher 12-550-15). The following antibodies were used: rabbit anti-IBA1 (1:1,000, Wako 019-19741), chicken anti-GFP (1:500, Fisher A10262), rabbit anti-mCherry (Fisher NBP2-25158), and mouse anti-CD68 (1:1,000, Fisher 14-0681-82). All staining was done on free floating tissue. First sections were placed in wells containing 10% Normal Goat Serum (Fisher 16210072), 1% Bovine Serum Albumin (Fisher BP1600-100), and 0.03% Triton X-100 (Sigma X100) in PBS for 1 hour at room temperature, then transferred to a primary antibody solution suspended in a staining buffer (1% BSA, 1% NGS, and 0.3% Triton X-100) on an orbital shaker at 4 °C. Following three washes in PBS, sections were incubated with AlexaFluor conjugated secondary antibodies, all at 1:600 dilution. Secondaries used were: Goat anti-Rabbit Alexa Fluor 568 (Life Technologies A11011), Goat anti chicken AlexaFluor 488 (Life Technologies A11039), Goat anti mouse Alex Fluor 647 (Fisher A21235). Hoescht 3342 (Fisher H3570) was used to stain nuclei, suspended at 1:1,000. All secondaries were suspended in the same staining buffer for 1 hour at room temperature. Sections were washed an additional three times, then mounted and coverslipped with Fluoromount-G (Southern Biotech 0100-01).

### Tissue imaging and processing

Tissue sections were imaged with the Zeiss Axioscan A2 Imager for 5x images of whole brain sections, and the ECHO Revolve Microscope for 20x magnification z-stacks used in the Sholl Analysis. Whole brain images were tiled together using the photomerge tool in Adobe Photoshop. All image filenames were hidden prior to analysis using the Fiji Blind Image Analysis Tool Plugin. Counts and CD68 intensities were analyzed via CellProfiler v 4.2.7. Area covered by the grafted cells was calculated using the Fiji Polygon selection tool to highlight the region colonized by the transplant, and densities were calculated by using the threshold 488/GFP channel in a selected region of interest and running the analyze particles feature (size = 0-0.1 inches, circularity 0.4 – 1). CD68 intensity was measured by creating masks of the 568/IBA1 channel using the analyze particles feature, and overlaying these ROIs on the 650/CD68 channel. Puncta which overlapped with IBA1+ signal was measured for intensity and normalized to the background intensity of each image. The Sholl analysis was processed by using the Sholl analysis plugin in the Imaris 3D image software (version 9.5.1) after using their automatic filament finding feature and adjusting the resulting filaments to match the cells visualized in the staining. All statistical analyses were two-way ANOVAs with Tukey correction performed on Graphpad Prism (version 10.4.1).

### Immunocytochemistry

Cells for analysis were placed in 12 well glass bottom plates (Cellvis, P12-1.5P) precoated with PDL. Prior to fixation, cells were washed three times with sterile 1X PBS, then placed in 4% PFA solution for 30 minutes. Wells were filled with blocking solution: 3 mg/mL normal goat serum (ThermoFisher 31872), 0.1mg/mL bovine serum albumin (Fisher BP9700100), 0.3% Triton X-100 in 1X PBS for 1 hour, followed by primary antibody in staining buffer (same as blocking solution, with 0.3 mg/mL goat serum) overnight at 4 °C. After 3 washes with 1X PBS, cells were treated with a secondary antibody solution in staining buffer before being placed under a coverslip within the well with Fluoromount-G. Cells were imaged using the REVOLVE microscope.

### Enzyme linked immunosorbent assay

For in vitro analysis, microglia were transduced overnight with the BDNF lentivirus as described previously in the methods. Following this, the microglia were treated with doxycycline dissolved in cell culture media at 100 nM for 72 hours. Media was then collected and used for BDNF ELISA analysis. Control cells were transduced with a GFP-lentivirus. For tissue ELISA, mice were euthanized prior to collecting tissue samples via a 1 mm biopsy punch (Fisher 12-460-402). Tissue samples were collected 0.5 mm posterior to the surgery site the location of which was measured using the stereotactic needle apparatus. Samples were immediately flash-frozen by submersion in liquid nitrogen. Prior to use, samples were thawed on ice for X min and placed in 200 µL of lysis buffer: 100 mM Tris (Fisher, BP152-5), pH 7.4, 150 mM NaCl (Sigma, S9888), 1 mM EGTA (R&D Systems, 2807/1G), 1 mM EDTA (Fisher, S311-500), 1% Triton X-100, 0.5% sodium deoxycholate (Sigma D6750), PMSF (R&D Systems, 448650), and a protease/phosphatase inhibitor cocktail (Fisher, PI78441). The tissue was then sonicated for 3 seconds on ice to ensure uniform homogenization. Samples were centrifuged at 15,000 g at 4 °C for 20 minutes. Supernatant was then collected and stored at -80 °C until used with the Quantikine BDNF Total ELISA kit (R&D Systems, DBNT00), as described by the manufacturer.

### Two-photon microscopy

Mice were pretreated with an intraperitoneal injection (IP) of dexamethasone (Hikma) at 1 mg/kg one hour prior to the start of surgery. Using the same surgical setup as for microglial transplantation, the mouse skull was scored using a scalpel and coated with Optibond (Omni Dental, 2342278) cured twice for 10 seconds using blue light. Following this, a ∼3 mm craniotomy was performed at -1.5 mm, -1 mm from Bregma, taking care to preserve the integrity of the meninges. A 3 mm coverslip (Warner, 64-0720) was affixed to the skull with cyanoacrylate adhesive. Mice were imaged before and after IP administration of 1 mg/kg lipopolysaccharide (Sigma, L4516-1MG), two weeks following coverslip placement. Mice were maintained on 1.5% isoflurane and a 37 °C heating pad throughout the imaging sessions on a two-photon microscope (Thorlabs Bergamo), equipped with a Coherent Vision S laser light source set to 920 nm and a 16x, 0.8 NA water immersion objective (Nikon). Microglia in the top 100 µm of the cortex were imaged with a step size of 0.5 µm every 10 minutes for one hour. Z-stacks were imported to Imaris 3D imaging software for analysis. Motility was analyzed using the Imaris (Oxford Instruments) automatic surface generation following thresholding and background subtraction of the images.

### Single-cell RNA sequencing

Six weeks after transplanting *Cx3Cr1^CreER^;Rosa-DTA;CSF1R ^fl/fl^* mice were given a lethal dose of isoflurane, and cells from one hemisphere were dissociated using the MACS Adult Neural Dissociation Kit (Miltenyi, 130-092-628) according to the manufacturer’s instructions. Briefly, cells were fixed using GEM-X Flex Sample Preparation v2 Kit (10X, 1000781), cDNA libraries were generated using the Chromium Fixed RNA Kit, Mouse Transcriptome (10x Genomics, 1000496), according to manufacturer’s instructions. Libraries were sent to Novogene for sequencing (NovaSeq X Plus Series PE150) with 400 million paired-end reads per library. Data was analyzed using 10X Cell Ranger software v8.0.1. Downstream analysis was performed using RStudio, Seurat v5[57], ScType[35], and Monocle3 Pseudotime[58]. Clustering was performed at 0.8 resolution, and microglia were identified using the ScType classification system before reclustering via UMAP at the same resolution and running the subsets through pseudotime analysis. All sequencing data used for this study are available under the accession code GSE301016.

### Western blot

Using the same supernatant generated for tissue ELISA (above), the sample aliquots were mixed at 1 µg/µL in LSDS Sample Buffer (Thermo, J61337AC) diluted in Nuclease Free Water (Fisher, AM9937). Samples were boiled at 90 °C for 10 minutes before loading, loaded at 10 µG per well into a SurePAGE Precast Gel 4-20% Bis-Tris (GenScript, M00657), and run at 120V for 2 hours. Protein was transferred to nitrocellulose membrane (Sigma, GE10600023) in transfer buffer: 2.9 g Tris Base (Sigma, 10708976001), 14.5 g glycine (Fisher, BP381-1), and 200 mL methanol (Fisher, A412-4) in 1 L Milli-Q water. The transfer was run at a constant 0.2 mA for 1 hour.

Membranes were blocked for 1 hour at room temperature with 5% BSA in TBS-T: 2.4 g Tris-HCl (Fisher, BP153-1), 5.6 g Tris Base, 8.8 g NaCl, 0.1 % Tween-20 (Sigma, P7949), pH 7.6 in Milli-Q water. Membranes were transferred to overnight primary antibody solution containing either: Rabbit anti-phospho-TrkB (Tyr706) (1:500, Fisher, PIPA5105012), rabbit anti-TrkB (1:1,000, Cell Signaling, P1045598), or mouse anti-GAPDH (1:1,000, Cell Signaling, 97166T) in the same blocking solution at 4 °C. After 3 washes in 1X TBS-T, membranes were soaked in secondary anti-rabbit or anti-mouse HRP-linked antibody (1:5000, Cell Signaling, 7074 and 7076, respectively) suspended in the same blocking solution for one hour at room temperature.

Following 3 washes with 1X TBS-T, membranes were visualized using a chemifluorescent imager (Azure Biosystems 600). Between different primary antibody stains, the gels were washed once for ten minutes in stripping buffer (0.15% glycine, 2% SDS, 1% Tween-20 in Nuclease Free Water at pH 2.2), followed by two ten-minute washes with 1X PBS, then two ten-minute washes with 1X TBS-T. Gels were then placed in blocking solution for additional primary staining following the same protocol. Gels were analyzed via ImageJ (1.54p), using the built in Gel Analyzer tool, ROIs were drawn to capture all visible bands. Reported values are all normalized to GAPDH values, all calculated as area occupied under the peaks on resulting “Plot Lanes” function call. All bands associated with a given protein were quantified, normalized to GAPDH expression, and averaged together.

## Supporting information

Hofland et al. Supplementary Figures

## Availability of Data and Materials

All scRNA-seq data is available at the Nacional Center for Biotechnology Information data repository under the accession code GSE301016. All code for the data analysis in this paper is openly accessible at https://github.com/GoncalvesLab/Hofland-et-al-microglia.

## Acknowledgements

We would like to thank the Flow Cytometry, Imaging, and Sequencing cores at the Albert Einstein College of Medicine.

## Online Supplemental Material

Supplemental figure 1 describes the culture purification protocol undertaken to generate microglial populations for transplantation. Supplemental figure 2 is the associated quantification of tissue used in the ELISA, scRNA, and two-photon experiments, along with the vector design of the BDNF lentivirus. Supplemental figure 3 illustrates additional sequencing analysis, including whole brain cell type breakdowns, variably expressed genes, and regression analyses across various sequenced microglial populations. Supplemental figure 4 demonstrates caCSF1R and mCherry lentiviral vector designs, along with in vitro quantification of vector activation in the presence of doxycycline.

---

<u>Supplemental figure 1. Primary microglial culture protocol:</u> (A) Schematic for microglial-like cell culture purification technique. (B, C) Representative images of microglia post-shake (B) and pre-shake (C). (D) Quantification of the percent of GFP+ cells in the well pre- and post-purification. N = 4 biological replicates. Analysis via unpaired t-test, **** = p < 0.0001.

<u>Supplemental figure 1. Primary microglial culture protocol:</u> (A) Schematic for microglial-like cell culture purification technique. (B, C) Representative images of microglia post-shake (B) and pre-shake (C). (D) Quantification of the percentage of GFP+ cells in the well pre- and post-purification. n = 4 biological replicates. Analysis via unpaired t-test, **** < p < 0.0001.

<u>Supplemental figure 2. IHC analysis of ELISA, two-photon, and scRNA tissue:</u> (A) Plasmid map for the BDNF lentivirus. (B, C, D) Representative images of the transplants for the ELISA, 2-photon, and scRNA sequencing experiments, respectively. (E) Quantifications of GFP+ cells in the relevant study areas. This was defined as throughout the brain for the scRNA sequencing experiment, and within 100 μm of the punch site/cover slip site on the ELISA and 2 photon tissue. (F) Representative traces from 20x images of scRNA microglial cells, along with Sholl analysis (G).

<u>Supplemental figure 3. Additional sequencing analysis:</u> (A) Full UMAP projection of whole brain samples, with clusters run through sctype classification prior to downstream analysis. All downstream work was performed in the “Microglial cells” subset. (B) Volcano plot of most differentially expressed genes amongst the microglial cell subset. (C) Feature Plots of relevant cellular markers for the cell clusters identified in Figure 5. (D) Feature plots of cell specific cell markers: P2ry12 (microglia), Aif1 (immune marker), Sox9 (astrocytes), Mki67 (proliferating cells), Plp1 (Dendritic cells), and S100a9 (Oligodendrocytes). All correlation plots were generated using differential expression gene lists relative to the E-MG data. (E) Correlation plot between Host and Transplanted cells from Figure 4B. (F) Correlation within the transplanted cells between those that express BDNF and those that do not. (G) Correlation between male and female microglial cells. n = 4 mice in both WT and Transplanted groups, 2 males and 2 females. (H) Correlation between T-MGs and R-MGLs from a separate data set[40].

<u>Supplemental figure 4. caCSF1R and mCherry infection strategies:</u> (A, B) Plasmid maps of the *caCSF1R* and *mCherry* control viruses. (C, D) Quantification of GFP+ cell density across various titers of the caCSF1R virus, both with and without doxycycline treatment. (E, F) Representative images of *CSF1R* cKO mice transplanted with mCherry control virus, both with and without doxycycline respectively. (G) Quantification of mCherry expression amongst GFP+ cells. (H) Representative images of mCherry and CaCSF1R transplants in CSF1R cKO mice treated with PLX for 1 week following 2 weeks of maintenance on tamoxifen. (I) Density of mCherry and caCSF1R infected GFP+/IBA1+, set relative to the GFP-/IBA1+ cell density within each individual brain to control for variable PLX response per animal. N = 4 animals per transplant, analysis performed via two-way ANOVA with Bonferroni correction

## References

1. Lewiecki, E.M., Biological therapy: chronicling 15 years of progress. Expert Opin Biol Ther, 2015. 15(5): p. 619–21.

2. Nunes, D., J.A. Loureiro, and M.C. Pereira, Drug Delivery Systems as a Strategy to Improve the Efficacy of FDA-Approved Alzheimer’s Drugs. Pharmaceutics, 2022. 14(11).

3. Pardridge, W.M., Blood-Brain Barrier and Delivery of Protein and Gene Therapeutics to Brain. Front Aging Neurosci, 2019. 11: p. 373.

4. Walker, F.R., et al., Dynamic structural remodelling of microglia in health and disease: a review of the models, the signals and the mechanisms. Brain Behav Immun, 2014. 37: p. 1–14.

5. Nimmerjahn, A., F. Kirchhoff, and F. Helmchen, Resting microglial cells are highly dynamic surveillants of brain parenchyma in vivo. Science, 2005. 308(5726): p. 1314–8.

6. Huang, Y., et al., Repopulated microglia are solely derived from the proliferation of residual microglia after acute depletion. Nat Neurosci, 2018. 21(4): p. 530–540.

7. Xu, Z., et al., Efficient Strategies for Microglia Replacement in the Central Nervous System. Cell Rep, 2020. 33(8): p. 108443.

8. Var, S.R., et al., Transplanting Microglia for Treating CNS Injuries and Neurological Diseases and Disorders, and Prospects for Generating Exogenic Microglia. Cell Transplant, 2023. 32: p. 9636897231171001.

9. Zhan, L., et al., Proximal recolonization by self-renewing microglia re-establishes microglial homeostasis in the adult mouse brain. PLoS Biol, 2019. 17(2): p. e3000134.

10. Lund, H., et al., Competitive repopulation of an empty microglial niche yields functionally distinct subsets of microglia-like cells. Nat Commun, 2018. 9(1): p. 4845.

11. Cronk, J.C., et al., Peripherally derived macrophages can engraft the brain independent of irradiation and maintain an identity distinct from microglia. J Exp Med, 2018. 215(6): p. 1627–1647.

12. Wilkinson, F.L., et al., Busulfan conditioning enhances engraftment of hematopoietic donor-derived cells in the brain compared with irradiation. Mol Ther, 2013. 21(4): p. 868–76.

13. Shibuya, Y., et al., Treatment of a genetic brain disease by CNS-wide microglia replacement. Sci Transl Med, 2022. 14(636): p. eabl9945.

14. Chen, D., et al., Brain-wide microglia replacement using a nonconditioning strategy ameliorates pathology in mouse models of neurological disorders. Sci Transl Med, 2025. 17(796): p. eads6111.

15. Chadarevian, J.P., et al., Harnessing human iPSC-microglia for CNS-wide delivery of disease-modifying proteins. Cell Stem Cell, 2025.

16. Hock, C., et al., Region-specific neurotrophin imbalances in Alzheimer disease: decreased levels of brain-derived neurotrophic factor and increased levels of nerve growth factor in hippocampus and cortical areas. Arch Neurol, 2000. 57(6): p. 846–51.

17. Gao, L., et al., Brain-derived neurotrophic factor in Alzheimer’s disease and its pharmaceutical potential. Transl Neurodegener, 2022. 11(1): p. 4.

18. Cardona, A.E., et al., Control of microglial neurotoxicity by the fractalkine receptor. Nat Neurosci, 2006. 9(7): p. 917–24.

19. Qin, K., et al., A much convenient and economical method to harvest a great number of microglia. Cell Mol Neurobiol, 2012. 32(1): p. 67–75.

20. Woolf, Z., et al., Isolation of adult mouse microglia using their in vitro adherent properties. STAR Protoc, 2021. 2(2): p. 100518.

21. Imai, Y., et al., A novel gene iba1 in the major histocompatibility complex class III region encoding an EF hand protein expressed in a monocytic lineage. Biochem Biophys Res Commun, 1996. 224(3): p. 855–62.

22. Kiani Shabestari, S., et al., Absence of microglia promotes diverse pathologies and early lethality in Alzheimer’s disease mice. Cell Rep, 2022. 39(11): p. 110961.

23. Elmore, M.R., et al., Colony-stimulating factor 1 receptor signaling is necessary for microglia viability, unmasking a microglia progenitor cell in the adult brain. Neuron, 2014. 82(2): p. 380–97.

24. Das, A., et al., Transcriptome sequencing reveals that LPS-triggered transcriptional responses in established microglia BV2 cell lines are poorly representative of primary microglia. J Neuroinflammation, 2016. 13(1): p. 182.

25. Hammond, T.R., et al., Single-Cell RNA Sequencing of Microglia throughout the Mouse Lifespan and in the Injured Brain Reveals Complex Cell-State Changes. Immunity, 2019. 50(1): p. 253–271 e6.

26. Cadiz, M.P., et al., Culture shock: microglial heterogeneity, activation, and disrupted single-cell microglial networks in vitro. Mol Neurodegener, 2022. 17(1): p. 26.

27. Wurm, J., et al., Microglia Development and Maturation and Its Implications for Induction of Microglia-Like Cells from Human iPSCs. Int J Mol Sci, 2021. 22(6).

28. Swanson, M.E.V., et al., Microglial CD68 and L-ferritin upregulation in response to phosphorylated-TDP-43 pathology in the amyotrophic lateral sclerosis brain. Acta Neuropathol Commun, 2023. 11(1): p. 69.

29. Morrison, H.W. and J.A. Filosa, A quantitative spatiotemporal analysis of microglia morphology during ischemic stroke and reperfusion. J Neuroinflammation, 2013. 10: p. 4.

30. Li, Q. and B.A. Barres, Microglia and macrophages in brain homeostasis and disease. Nat Rev Immunol, 2018. 18(4): p. 225–242.

31. de Pins, B., et al., Conditional BDNF Delivery from Astrocytes Rescues Memory Deficits, Spine Density, and Synaptic Properties in the 5xFAD Mouse Model of Alzheimer Disease. J Neurosci, 2019. 39(13): p. 2441–2458.

32. Andrade-Moraes, C.H., et al., Cell number changes in Alzheimer’s disease relate to dementia, not to plaques and tangles. Brain, 2013. 136(Pt 12): p. 3738–52.

33. Honey, D., et al., Analysis of microglial BDNF function and expression in the motor cortex. Front Cell Neurosci, 2022. 16: p. 961276.

34. Das, A.T., L. Tenenbaum, and B. Berkhout, Tet-On Systems For Doxycycline-inducible Gene Expression. Curr Gene Ther, 2016. 16(3): p. 156–67.

35. Ianevski, A., A.K. Giri, and T. Aittokallio, Fully-automated and ultra-fast cell-type identification using specific marker combinations from single-cell transcriptomic data. Nat Commun, 2022. 13(1): p. 1246.

36. Ruan, C. and W. Elyaman, A New Understanding of TMEM119 as a Marker of Microglia. Front Cell Neurosci, 2022. 16: p. 902372.

37. Matcovitch-Natan, O., et al., Microglia development follows a stepwise program to regulate brain homeostasis. Science, 2016. 353(6301): p. aad8670.

38. Colella, P., et al., CNS-wide repopulation by hematopoietic-derived microglia-like cells corrects progranulin deficiency in mice. Nat Commun, 2024. 15(1): p. 5654.

39. Gerrits, E., et al., Transcriptional profiling of microglia; current state of the art and future perspectives. Glia, 2020. 68(4): p. 740–755.

40. Shemer, A., et al., Engrafted parenchymal brain macrophages differ from microglia in transcriptome, chromatin landscape and response to challenge. Nat Commun, 2018. 9(1): p. 5206.

41. Elmore, M.R., et al., Characterizing newly repopulated microglia in the adult mouse: impacts on animal behavior, cell morphology, and neuroinflammation. PLoS One, 2015. 10(4): p. e0122912.

42. Roussel, M.F., et al., A point mutation in the extracellular domain of the human CSF-1 receptor (c-fms proto-oncogene product) activates its transforming potential. Cell, 1988. 55(6): p. 979–88.

43. Morley, G.M., et al., Cell specific transformation by c-fms activating loop mutations is attributable to constitutive receptor degradation. Oncogene, 1999. 18(20): p. 3076–84.

44. Wang, Y., et al., TrkB/BDNF signaling pathway and its small molecular agonists in CNS injury. Life Sci, 2024. 336: p. 122282.

45. Yoo, Y., et al., A cell therapy approach to restore microglial Trem2 function in a mouse model of Alzheimer’s disease. Cell Stem Cell, 2023. 30(8): p. 1043–1053 e6.

46. Vay, S.U., et al., The plasticity of primary microglia and their multifaceted effects on endogenous neural stem cells in vitro and in vivo. J Neuroinflammation, 2018. 15(1): p. 226.

47. Svoboda, D.S., et al., Human iPSC-derived microglia assume a primary microglia-like state after transplantation into the neonatal mouse brain. Proc Natl Acad Sci U S A, 2019. 116(50): p. 25293–25303.

48. Abud, E.M., et al., iPSC-Derived Human Microglia-like Cells to Study Neurological Diseases. Neuron, 2017. 94(2): p. 278–293 e9.

49. Kennedy, D.W. and J.L. Abkowitz, Kinetics of central nervous system microglial and macrophage engraftment: analysis using a transgenic bone marrow transplantation model. Blood, 1997. 90(3): p. 986–93.

50. Rivera, V.M., et al., Long-term regulated expression of growth hormone in mice after intramuscular gene transfer. Proc Natl Acad Sci U S A, 1999. 96(15): p. 8657–62.

51. Chadarevian, J.P., et al., Engineering an inhibitor-resistant human CSF1R variant for microglia replacement. J Exp Med, 2023. 220(3).

52. Spangenberg, E., et al., Sustained microglial depletion with CSF1R inhibitor impairs parenchymal plaque development in an Alzheimer’s disease model. Nat Commun, 2019. 10(1): p. 3758.

53. Bosch, A.J.T., et al., CSF1R inhibition with PLX5622 affects multiple immune cell compartments and induces tissue-specific metabolic effects in lean mice. Diabetologia, 2023. 66(12): p. 2292–2306.

54. Varvel, N.H., et al., Replacement of brain-resident myeloid cells does not alter cerebral amyloid-beta deposition in mouse models of Alzheimer’s disease. J Exp Med, 2015. 212(11): p. 1803–9.

55. Chitu, V. and E.R. Stanley, Regulation of Embryonic and Postnatal Development by the CSF-1 Receptor. Curr Top Dev Biol, 2017. 123: p. 229–275.

56. Le, L.H.D., et al., The microglial response to inhibition of Colony-stimulating-factor-1 receptor by PLX3397 differs by sex in adult mice. Cell Rep, 2025. 44(1): p. 115176.

57. Hao, Y., et al., Dictionary learning for integrative, multimodal and scalable single-cell analysis. Nat Biotechnol, 2024. 42(2): p. 293–304.

58. Qiu, X., et al., Reversed graph embedding resolves complex single-cell trajectories. Nat Methods, 2017. 14(10): p. 979–982.

