## Supplementary figures and images for "Controlled Delivery of a Neurotrophic Factor in the Adult Mouse Brain Using Engineered Microglia"

### Hofland et al. Supplementary Figures

#### Supplemental figure 1

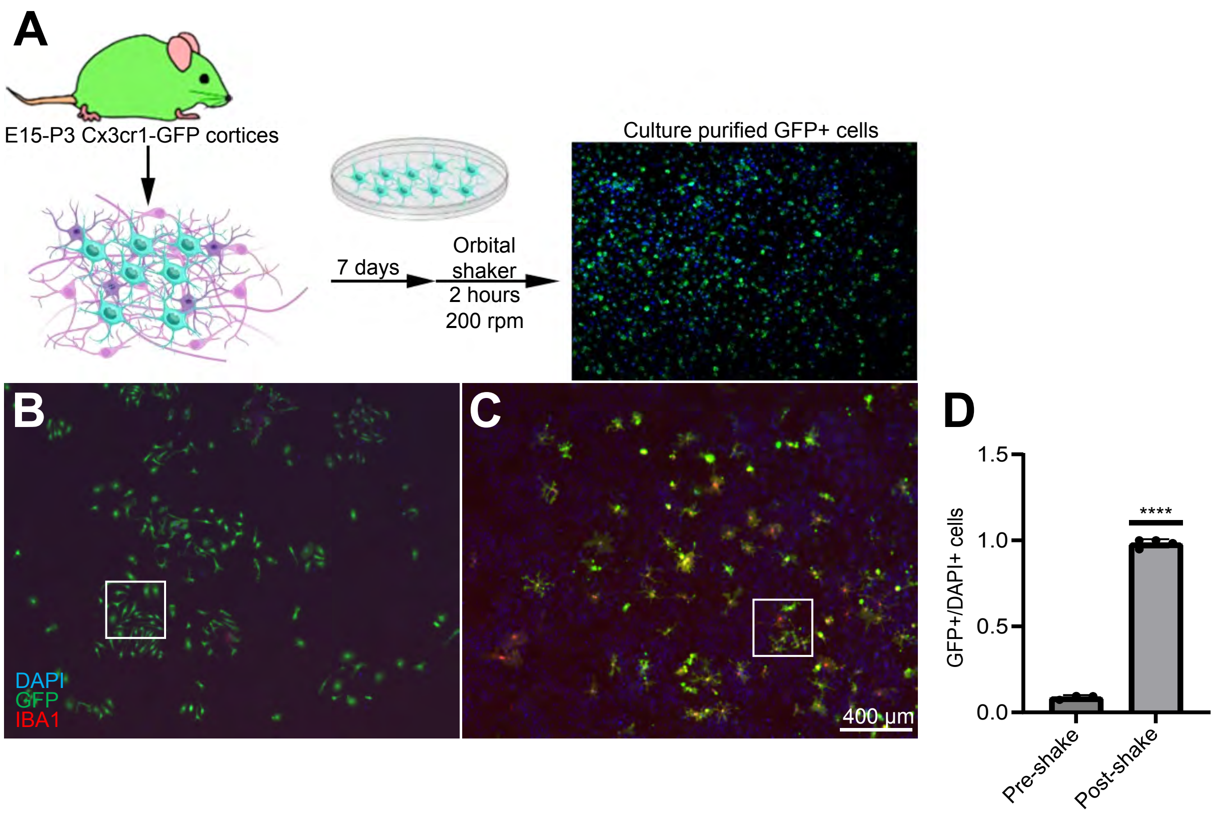

#### Supplemental figure 2

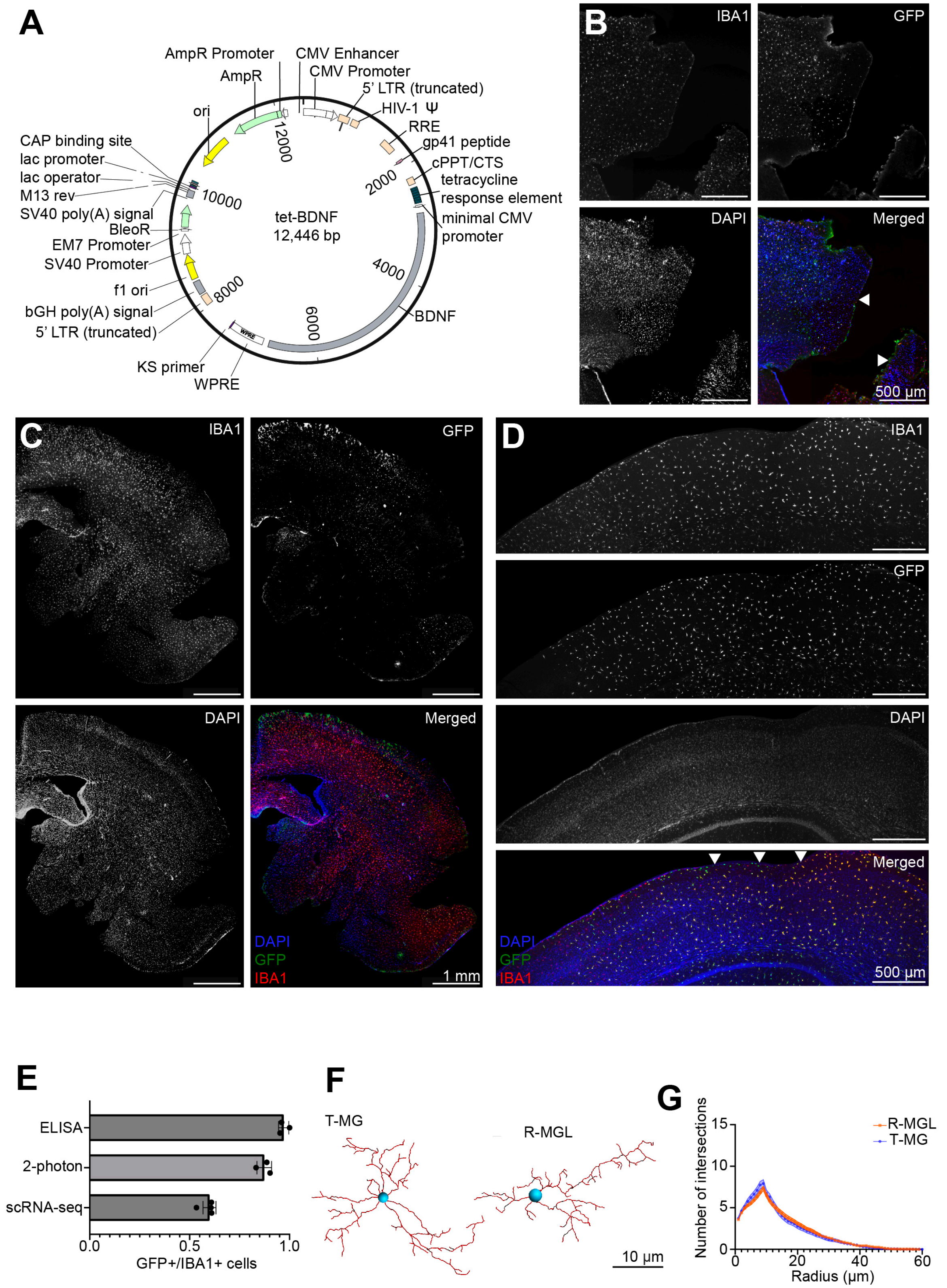

#### Supplemental figure 3

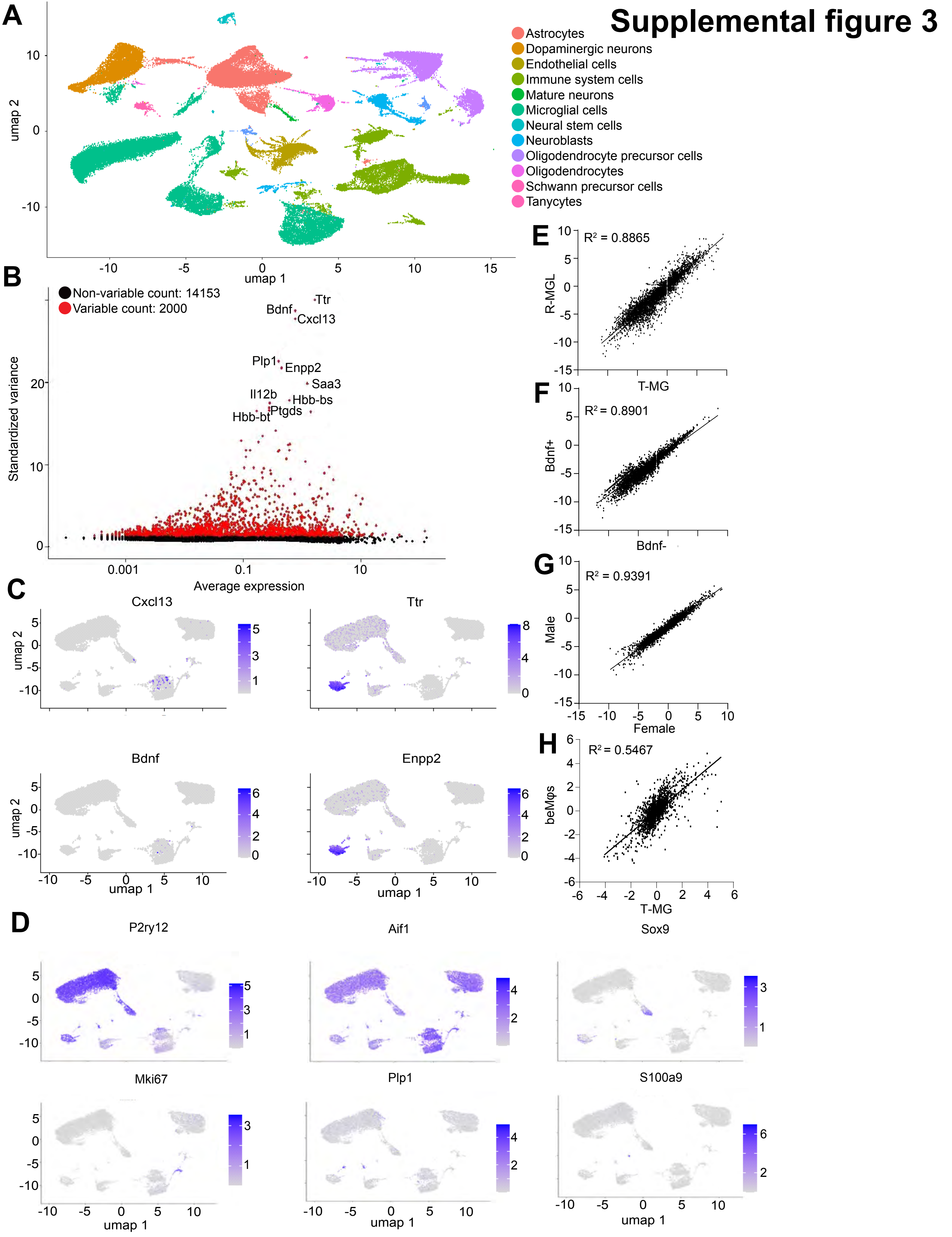

#### Supplemental figure 4

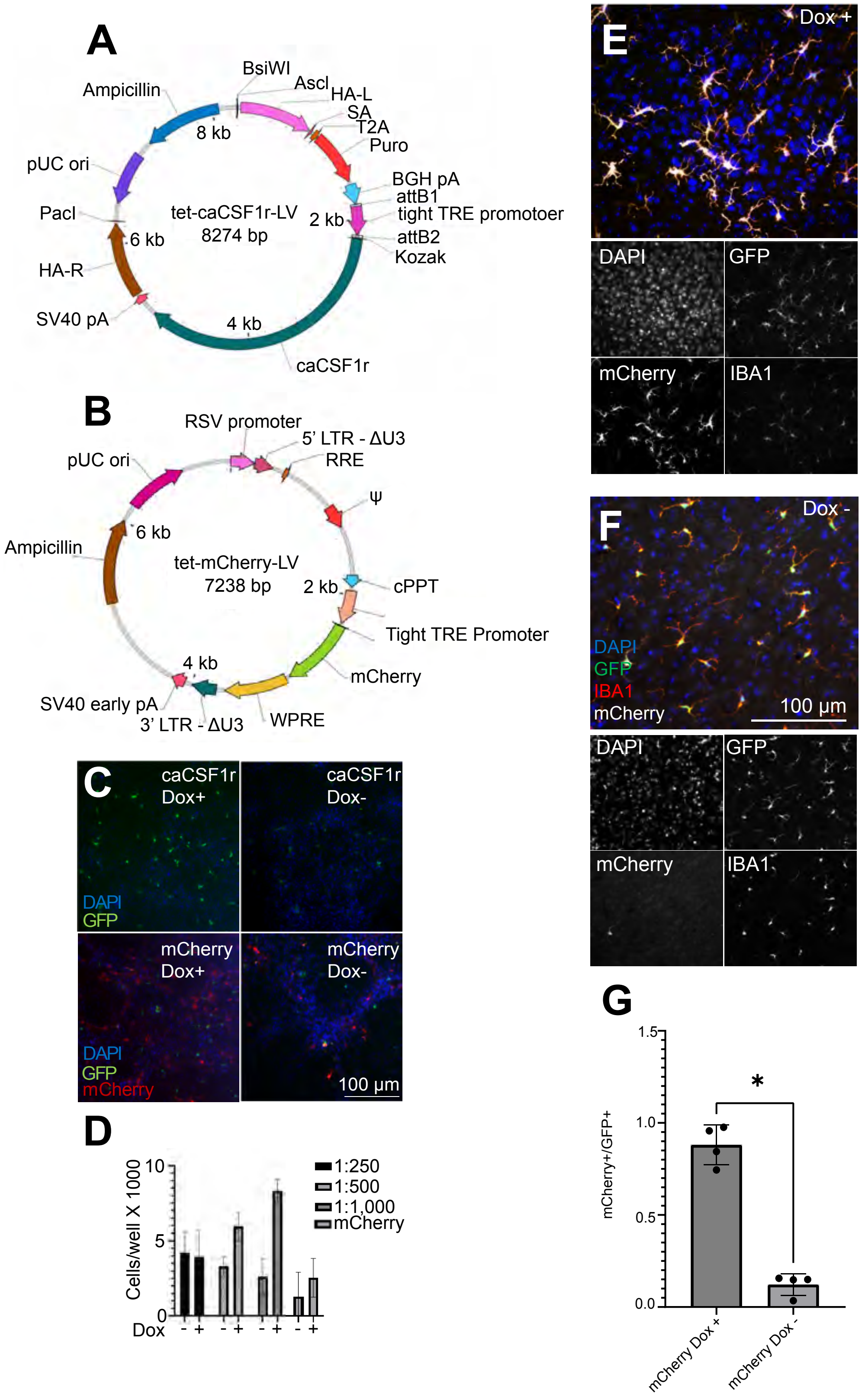
